# Comparative genomics reveals potential mechanisms of invasion in *Phragmites australis* (common reed)

**DOI:** 10.64898/2026.06.12.731924

**Authors:** Samadhi Wimalagunasekara, Richard S. Garcia, Thu T. Nguyen, Pramod Pantha, Guannan Wang, Dong-Ha Oh, Wesley A. Bickford, Kurt P. Kowalski, Keith Clay, Maheshi Dassanayake

## Abstract

Biological invasions are transforming ecosystems worldwide, yet the genomic bases enabling certain species to dominate new environments remain poorly understood. *Phragmites australis*, a widespread wetland grass with invasive and native subspecies co-occurring in North America, provides a powerful system to investigate genomic mechanisms of invasiveness. We generated independent chromosome-scale genome assemblies for invasive *P. australis ssp. australis* and co-occurring native *ssp. americanus* and used comparative genomic and transcriptomic analyses to identify lineage-specific innovations associated with invasive success. The invasive subspecies exhibits genomic novelties through functionally-biased single-copy orthologs, intronless genes, and subgenome expression asymmetry, along with a stress-ready basal transcriptome relative to the native subspecies. Following the removal of aboveground shoots (“cutback”), which measures the ability to recover from damage, the invasive subspecies undergoes stronger transcriptional reprogramming, increased shoot production, and higher biomass accumulation compared to the native. It also displays expansion of gene families and coordinately expressed gene modules that support resource mobilization, growth responses to light, and stress tolerance. Beyond *Phragmites*, comparative analyses across multiple grass genomes, including eight invasive species with related non-invasive species, revealed repeated expansion of gene families associated with abiotic stress tolerance and developmental regulation, suggesting convergent adaptive strategies in the grass family for invasive success. Together, these results demonstrate genomic architecture linked to invasion success and highlight potential targets for managing invasive grasses.

## Introduction

Invasive plants are defined by their aggressive growth habits and capacity to cause ecosystem damage when introduced into new environments. They outcompete native species and cause massive environmental, economic, and human health impacts with estimated costs rising each year (Diagne et al., 2021). Charles Elton articulated the foundational concept that ecological imbalances caused by invasive species can transform ecosystems and cause severe ecological disruption (Elton, 1958). The success of invasive plants is rooted in genetically-based life histories characterized by selfing and vegetative propagation, high seed production and rapid growth and broad environmental tolerance (Baker and Stebbins, 1965). Despite these early insights linking genetics to invasion success, most subsequent research on invasive plants has focused on population-level dynamics of individual species in their native vs. invaded ranges or on ecological comparisons between invaded vs. non-invaded plant communities. By contrast the genomic mechanisms underlying invasive ability remain poorly understood (Hodgins et al., 2025).

Invasive plants must adapt to novel biotic and abiotic pressures to establish and thrive in new environments. Key traits include robust growth, efficient resource use, resilience to stress, and successful reproduction despite limited genetic diversity resulting from founder effects (Baker and Stebbins, 1965; Liao et al., 2020). We do not know which genomic features or transcriptional responses enable rapid growth, stress tolerance, and regrowth following damage compared to closely related, non-invasive populations. In addition, the lack of comparative genomic analyses across diverse, unrelated invasive lineages makes it difficult to identify which genes or genetic networks are consistently divergent from related, non-invasive populations or species. Addressing these gaps could not only advance our understanding of genomic mechanisms of invasiveness but could also enable targeted interventions to mitigate impacts of invasive plants under changing environmental conditions.

*Phragmites australis* (Cav.) Trin. ex Steud. (common reed) is a tall wetland grass (Poaceae, tribe Arundineae) with an extensive network of rhizomes and stolons that supports extensive clonal growth (Packer et al., 2017). New populations are established by seed or vegetative fragments. The species is self-compatible, but seed set is higher in larger, more diverse populations (Kettenring et al., 2011; Lambert and Casagrande, 2007). In North America, *P. australis* includes three subspecies that vary in their degree of invasiveness, making it an ideal model for invasive ability given its widespread distribution and ecological impacts across diverse habitats (Christenhusz and Fay, 2024; Cui et al., 2025; Oh et al., 2022; Saltonstall, 2002; Wang et al., 2024b, 2024a). The invasive subspecies *australis* and the non-invasive subspecies *americanus* co-occur widely in the Laurentian Great Lakes region and exhibit variation within and between populations (Packer et al., 2017), with the subspecies *berlandieri* found along the US Gulf Coast with minimal overlap with the other two subspecies (Saltonstall and Hauber, 2007). The two subspecies examined here share the same whole genome duplication event 30 – 40 million years ago (Oh et al., 2022; Wang et al., 2024b). Whole-genome and tandem duplication events create genetic redundancy that can fuel rapid evolutionary innovation (Panchy et al., 2016; Soltis and Soltis, 2016). Preferential retention of genes following whole genome duplication and lineage specific tandem duplication events appeared generally enriched for regulatory genes and maintaining DNA integrity in the invasive *P. australis* lineage (Oh et al., 2022; Wang et al., 2024b). Studies on *P. australis* continue to provide insights into mechanisms driving invasions and inform management strategies for invasive species (Allen et al., 2020; Bickford et al., 2022; Ji et al., 2025; Kowalski et al., 2015; Meyerson et al., 2025; Tao et al., 2023; Wang et al., 2025).

*P. australis* occurs primarily as an allotetraploid in North America, while exhibiting a broader range of ploidy levels worldwide; however, a direct link between polyploidization and invasiveness was not established (Wang et al., 2024a). Available genomic resources for subspecies *australis* and *americanu*s (Oh et al., 2022; Wang et al., 2024b) do not provide independent chromosomal assemblies generated using comparable long-read sequencing, which would allow a direct comparison of reference genomes for both invasive and native subspecies coexisting in similar geographic regions. Because *P. australis* phenotypes collected from geographically distinct regions are unreliable indicators of genome size, ploidy level, or lineage-specific traits (Wang et al., 2024a), high quality genomes for both subspecies co-occurring in the same sites are essential to accurately identify their genomic differences potentially driving invasiveness in subspecies *australis*. Hybridization between native and invasive *P. australis* lineages in North America appears to be rare, and when identified, hybrids showed lower seed germination rates, while backcrossing with parental lineages appeared to be even rarer or remains poorly documented (Williams et al., 2019; Wu et al., 2015). Therefore, co-occurring stands of native and invasive *P. australis* subspecies are generally assumed to remain relatively genetically isolated.

In this study, we generated chromosome-level reference genomes for two *Phragmites australis* subspecies (one invasive and one non-invasive) that co-occur in North America. These high-quality assemblies enabled direct comparison of genomic traits that have evolved since the most recent shared whole-genome duplication event in this species. Continuing our comparative analyses, we examined transcriptomic responses to aboveground biomass removal (cutback) in multiple populations of both invasive and native *P. australis* subspecies. We then identified shared and divergent molecular responses, preadapted traits specific to each subspecies, and transcriptional changes uniquely associated with increased invasive capabilities. We extended this research to compare these genomes to all other available high-quality invasive grass species genomes along with those of related non-invasive grasses to identify emergent genomic features associated with invasiveness in the grass family. Our findings collectively highlight shared genomic features among invasive grasses, connect gene expression variation with traits that could promote invasiveness, and provide key genomic resources for developing management strategies against invasive *P. australis* and other invasive grasses.

## Results

### Independent chromosome-level reference genomes for invasive and native *Phragmites australis* in North America

#### Genome assembly

We assembled two independent, high-quality reference genomes representing the invasive and native subspecies of *Phragmites australis*, where the invasive European lineage is widespread across North America and is competing with the native lineage at multiple sites in the Laurentian Great Lakes region (Figure 1a, b and S1a, c) (Saltonstall, 2002). We generated PacBio continuous long-read (CLR) data, producing 224.5 Gb across 8.5 million reads for the native reference sample and 253 Gb across 7.2 million reads for the invasive reference sample. Each assembly was further scaffolded with Omni-C link libraries sequenced to yield paired-end Illumina sequences of 53.8 Gb for the native and 46.2 Gb for the invasive sample. Following gap filling, artifact removal, scaffolding, chromosome assignment, and subgenome phasing, we obtained genome assemblies of approximately 951 Mb for *P. australis ssp. americanus* (native) and 865 Mb for *P. australis ssp. australis* (invasive), which were used as reference genomes in all subsequent comparative analyses (Table 1). The genome files are available from NCBI under bioproject PRJNA705976 and Supplementary Files S1 and S2. The largest 24 scaffolds in each assembly correspond to pseudomolecules representing the 24 chromosomes (Fig. 1b).

**Figure 1.**
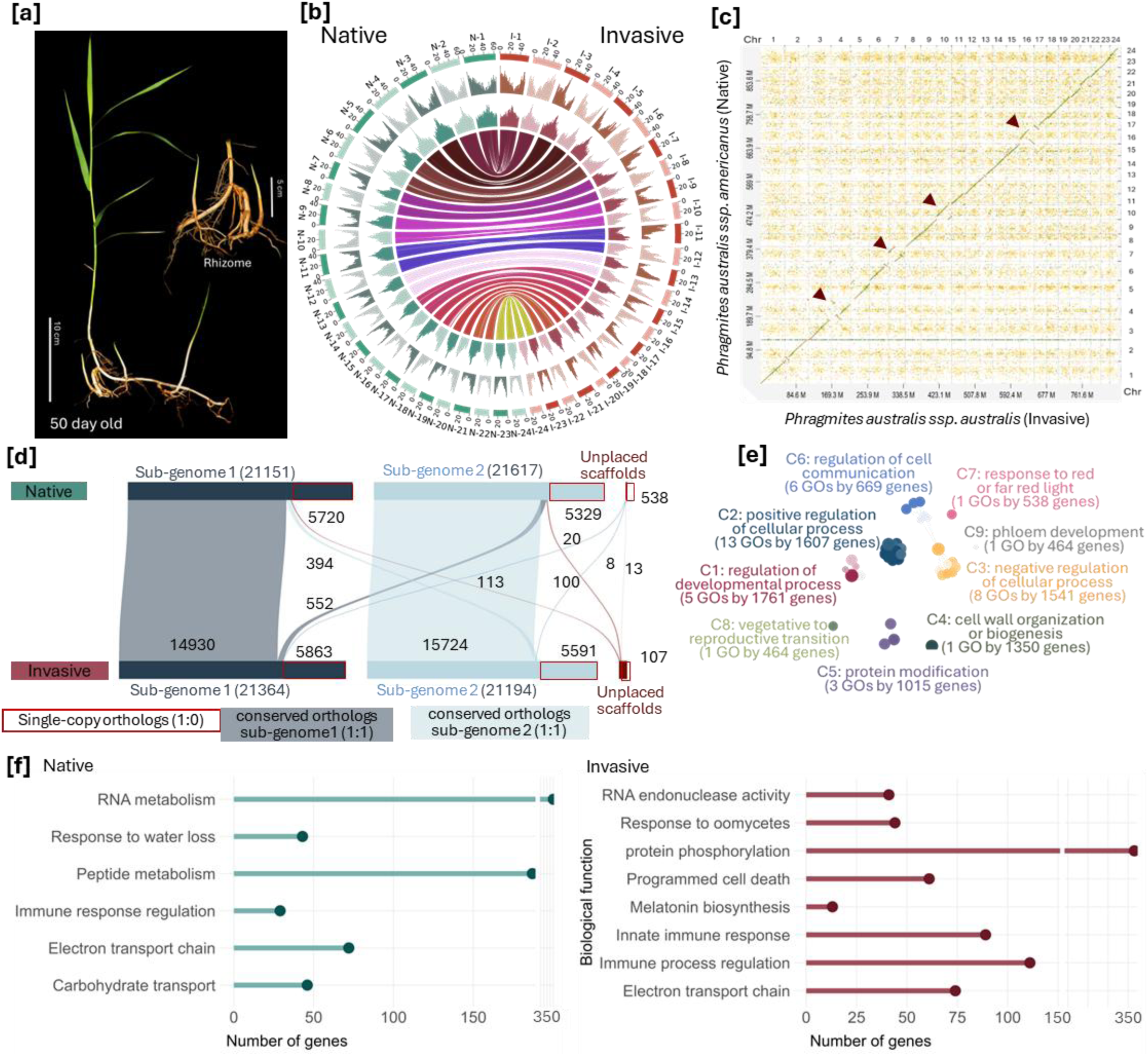
Genomes of invasive *Phragmites australis ssp. australis* and the native *P. australis ssp. americanus* are highly syntenic, showing conserved chromosome organization and similar subgenome architecture. [a] A plantlet of invasive *P. australis*. [b] *P. australis* genomes aligned to show chromosome-level synteny. Outer ring – annotated chromosomes N1-24 (green) for native and I1-14 (maroon) for invasive; chromosome pairs assigned to subgenomes (alternating dark and light blocks); odd numbered chromosomes – Subgenome1; even numbered chromosomes – Subgenome2; scale represents the chromosome size in Mb. Middle ring – gene density (per Mb). Inner ring – LTR density (per Mb). Innermost circle – colored lines between chromosomes show macrosynteny between genomes. [c] Whole genome alignment between invasive and native reference genomes shows high collinearity. Examples of interruptions to collinearity due to segmental duplications, deletions, and inversions are shown by brown arrow heads. [d] Gene composition and ortholog distribution between invasive and native subgenomes. Shaded regions represent syntenic 1:1 orthologs while lines represent translocated 1:1 orthologs between the two genomes. Red outlined boxes without any connecting lines represent single-copy orthologs (1:0), where a gene in one genome does not have an identifiable ortholog in the other genome. [e] Functional summary representation for conserved orthologs. [f] Functional summary for single-copy orthologs in native and invasive subspecies. All displayed functional summaries exclude overly broad (>2,000 genes) or very narrow (<10 genes) functional categories, while retaining non-redundant representative processes as much as possible. All enriched functions have adj p-values <0.05 with false discovery rate correction applied.

**Table 1.**
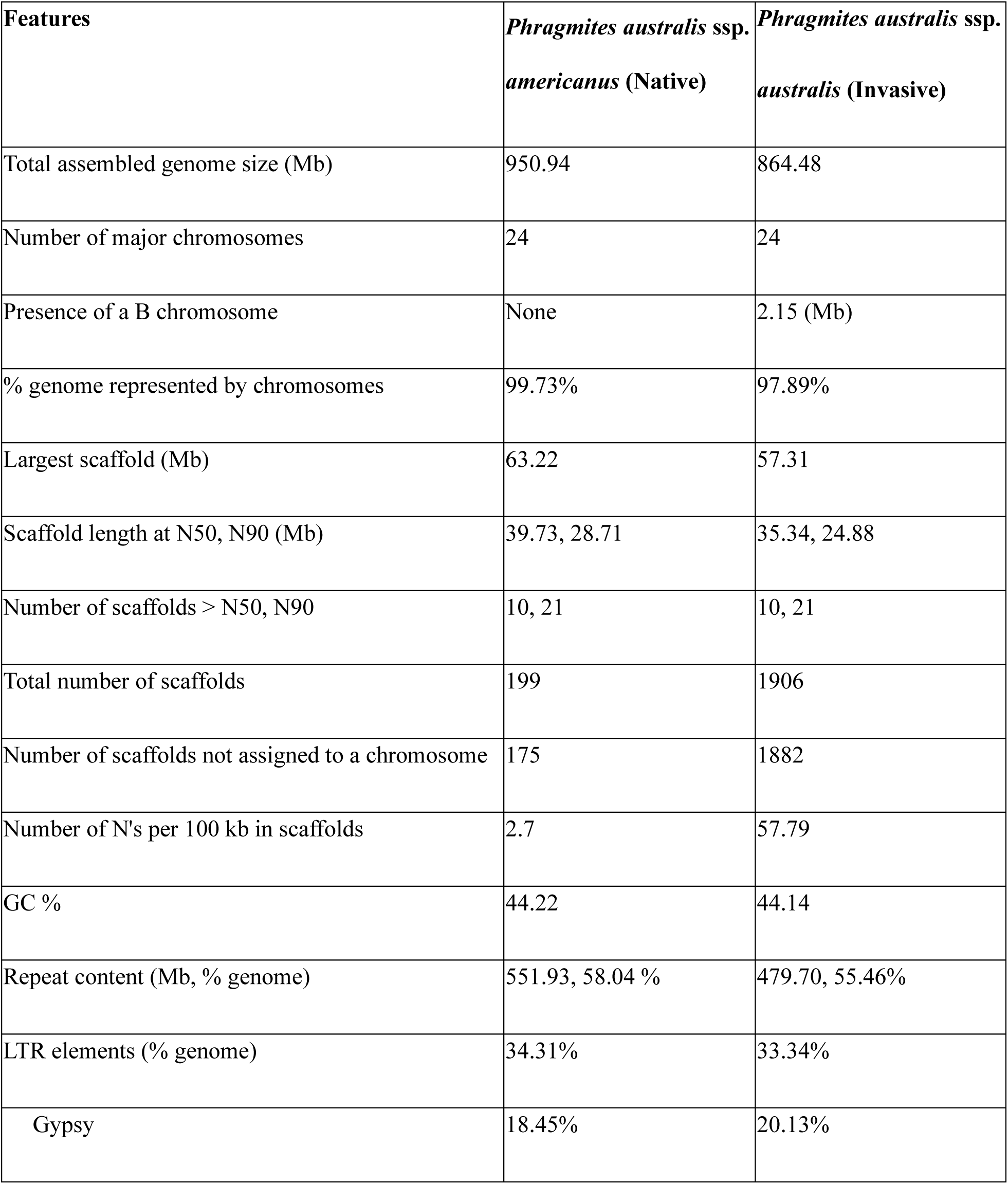
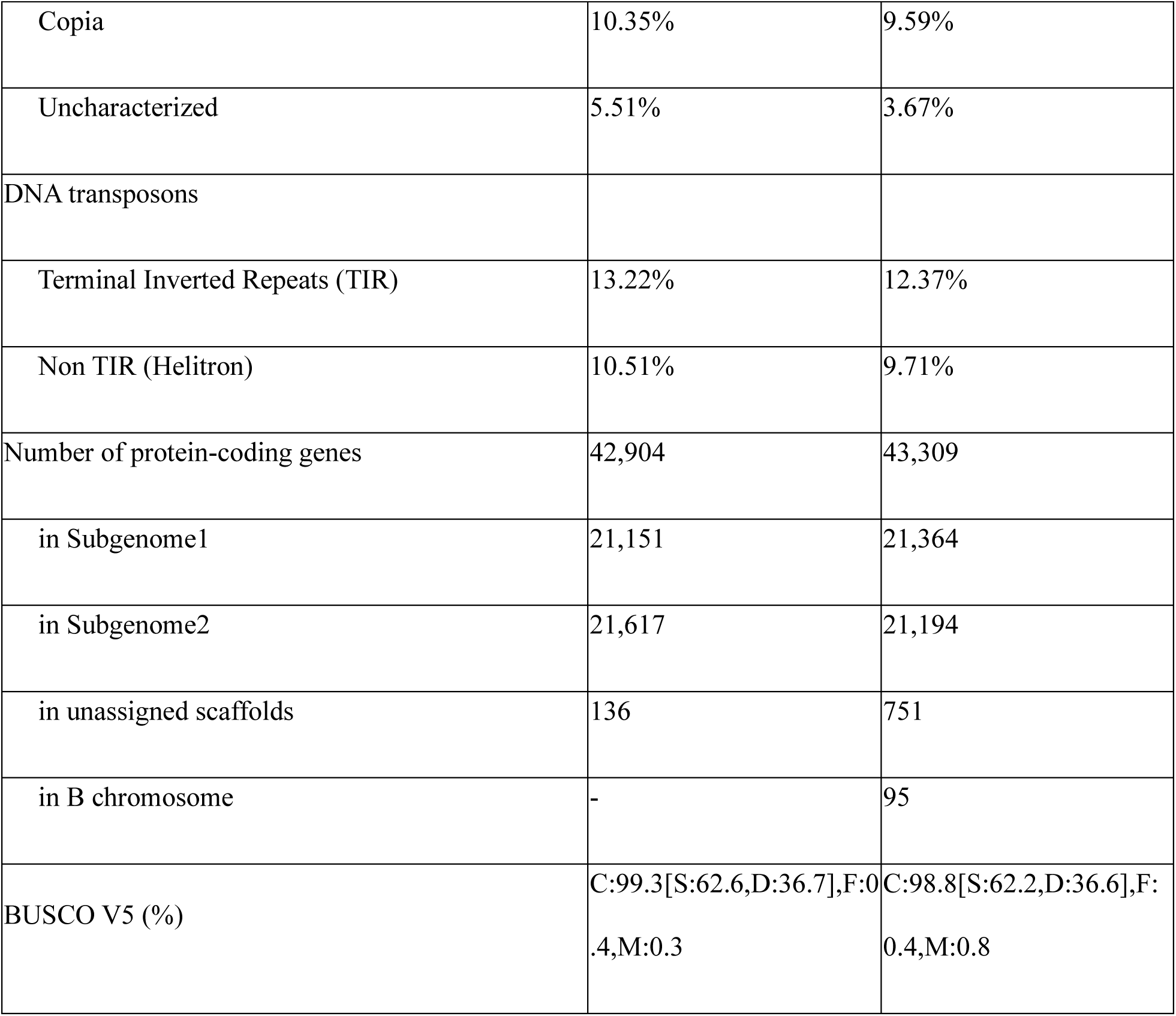
Summary of the genome assembly and structural annotation.

Only 0.3% (native) and 2% (invasive) of the genome was assigned to smaller unplaced scaffolds (Table 1, Supplementary File S3 and S4). The assembly sizes and the chromosome counts are consistent with the allotetraploid *P. australis* genomes previously reported (Christenhusz and Fay, 2024; Cui et al., 2025; Oh et al., 2022; Wang et al., 2024b, 2024a). While the two genomes are highly similar at the chromosome-level (macrosynteny) with broadly preserved collinearity (Fig. 1b-d), they diverge at finer scales (microsynteny) due to localized inversions, duplications, and deletions that impact gene-level organization (e.g., regions within chromosomes 4, 7, 10, and 17; Fig. 1c).

#### Genome architecture & synteny

Over 80% of each genome contains duplicated genomic blocks corresponding to the other (Figure S2a), consistent with tetraploid descendants of a shared whole-genome duplication event in the *P. australis* lineage. The presence of two subgenomes was also evident, as two chromosomes from each *P. australis* genome showed the highest similarity to a single rice chromosome (Fig. S2b). Homeologous chromosomes closely resembled their corresponding homeologs in the other genome (Fig. S2c). These homeologs had distinct repeat structures (Table S1), making it possible to differentiate the ancestral parental chromosome sets resulting in two subgenomes. Therefore, the 24 chromosomes in each reference assembly were assigned to two subgenomes, each consisting of 12 chromosomes, with even-numbered chromosomes representing subgenome 1 and odd-numbered chromosomes representing subgenome 2 (Fig. 1b). Both subgenomes show preserved collinearity between the native and invasive genomes across all main chromosomes (Fig. 1d, S2c). Genomic comparisons between rice and the *P. australis* genomes revealed extensive genomic rearrangements. At the same time, each *P. australis* genome also showed distinct rearrangements relative to the rice reference (Fig. S2d). The invasive reference genome is approximately 10% smaller than the native genome (Table 1), suggesting that the size difference primarily reflects cumulative structural rearrangements between the lineages.

#### B chromosome

Eleven unplaced scaffolds in the invasive genome showed high similarity to a previously reported B chromosome in *P. australis* from China (Cui et al., 2025) (Figure S3a). No comparable alignments were detected among scaffolds in the native genome. This supports reports of a single-copy B chromosome in invasive genotypes, but is absent in the native lineages (Wang et al., 2024b). Therefore, we combined the relevant scaffolds into a meta-scaffold and annotated it as a B chromosome following a reference guided assembly to the published B chromosome (Supplementary File S5). The newly-identified B chromosome in the invasive genome here is significantly smaller and contains more diverged sequences than the previously reported B chromosomes for *P. australis* (Cui et al., 2025; Wang et al., 2024b) (Fig. S3b).

#### Gene composition

Both genomes show a similar genic-to-non-genic composition (∼5% genic), with repeats comprising ∼55% of the genome with the native genome containing slightly more repeats than the invasive genome (Figure S4a). Gypsy and Copia long terminal repeat retrotransposons (LTRs) account for the majority of repeats (Table S1). Although reported LTR content varies among *P. australis* genomes, our results are consistent with the general pattern of LTR enrichment in pericentromeric regions and gene content concentrated toward chromosome ends (Fig 1b). We annotated ∼43,000 protein-coding genes for each genome (Table 1, Table S2, and Fig. 1d). Gene models were supported by *ab initio* predictions, conservation with proteins from other plants, and RNA-seq data generated in this study (Fig. S1b) and previously published (Oh et al., 2022). The protein coding sequences are available in Supplementary File S6 and 7. Currently annotated non-coding genes account for ∼0.05% in each genome. Among these, rRNA sequences represent the largest fraction, followed by snoRNAs and tRNAs (Fig. S4b). However, considering the number of distinct non-coding RNA types, more small non-coding RNAs and tRNAs were annotated than other classes (Fig. S4c). Among functionally annotated protein-coding genes representing conserved orthologs (1:1 copy) between the invasive and native genomes, core metabolic, cellular, and reproductive processes are predominant (Fig. 1e, Table S3.1). There has been extensive gene fractionation following their allopolyploid origin and subsequent diploidization in *P. australis*. As a result, ∼25% of genes in each genome were retained as single-copy orthologs where a gene in one genome did not have an identifiable ortholog in the other genome (Fig. 1d). By contrast, single-copy orthologs were enriched for functions related to biotic stress responses in *P. australis*, with the invasive genome containing a higher number of genes associated with pathogen responses than the native genome (Fig. 1f, Table S3.2). BUSCO assessments (v5, embryophyte dataset (Simão et al., 2015)) indicated high completeness, with 99.3% of single-copy orthologs recovered in the native genome and 98.8% in the invasive genome (Table 1). These *P. australis* assemblies are among the most complete and highly contiguous grass genomes available for comparative analysis.

### A stress-ready transcriptome distinguishes invasive from native subspecies

Mechanical removal of aboveground tissues (referred to as cutback or the “cut” treatment) is a widely used method to control invasive *P. australis* (Fig. S1d) and commonly occurs in grasses subjected to grazing or mowing. To evaluate transcriptomic responses, to cutback, we sampled six populations representing invasive and native subspecies co-occurring in the Great Lakes region (Fig. S1b). As each location with invasive or native genotypes, 28–32 shoot and rhizome samples were collected and regrown in a greenhouse to establish control (uncut) and treatment (cut) groups. This design comprised 24 groups (two subspecies, two tissues (shoot and rhizome), two treatments (uncut and cut), and three genotypes) represented by six to eight biological replicates per group, resulting in 186 RNA-seq libraries (Table S4). Our goal was to identify transcriptomic signatures distinguishing co-occurring invasive and native subspecies beyond genotype-level variation.

#### Global transcriptome profiles

Mean transcript expression decreased in all cutback samples, likely reflecting a global transcriptional response to loss of aboveground tissues. Transcriptomic differences between invasive and native subspecies were greater than variation among genotypes within each group (Figure 2a,b). Although rhizomes were not directly damaged by cutback, they showed greater transcriptional divergence before and after treatment than newly developed shoots relative to controls (Fig. 2a,b).

**Figure 2.**
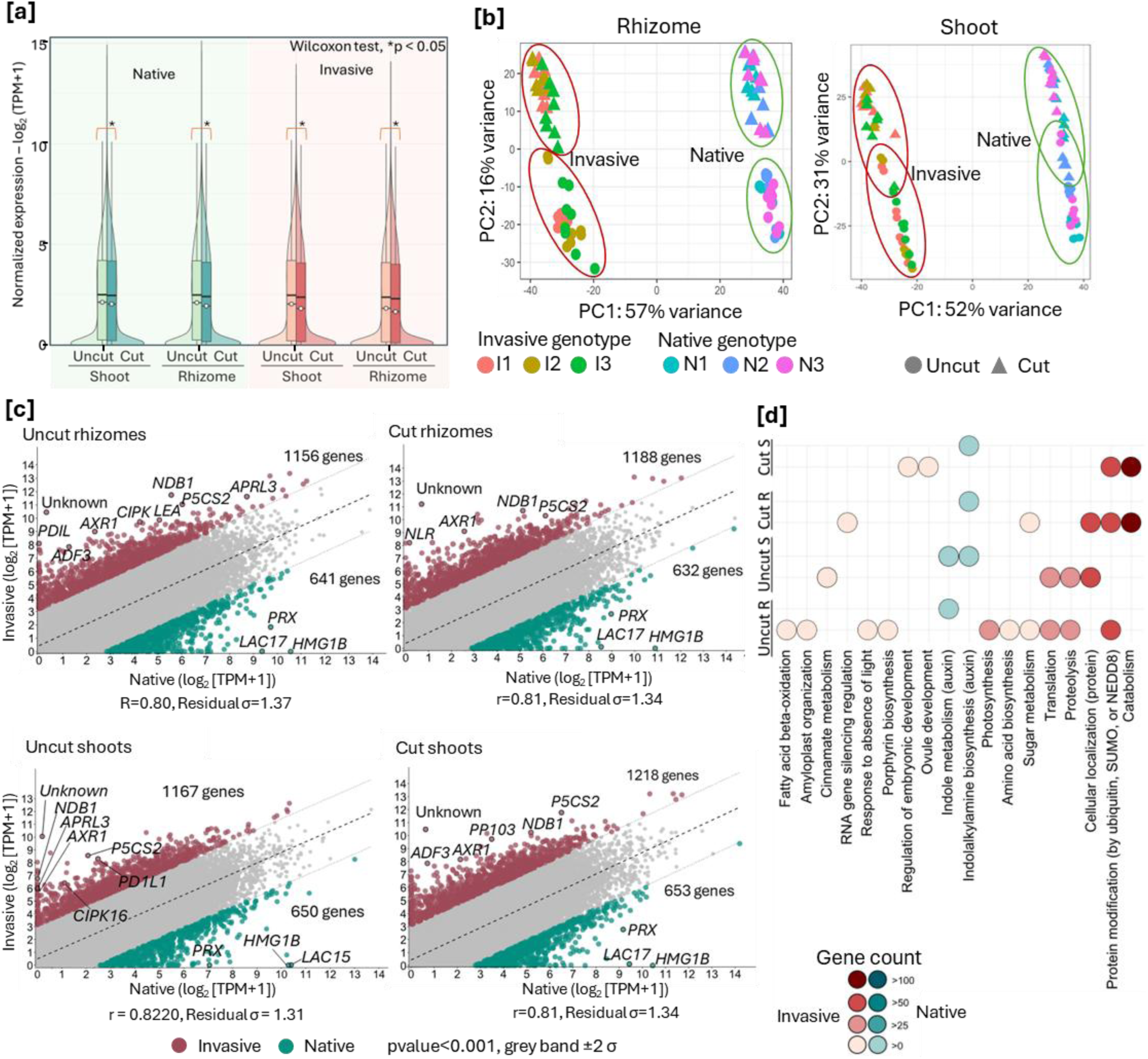
Distinct transcriptional states between cut and uncut samples include elevated basal expression that primes invasive transcriptomes for stress resilience and growth. [a] Total transcriptome expression profile before and after cutback. Mean expression is significantly lower for cut tissues compared to uncut tissues (Wilcoxon test, *p ≤ 0.05). Mean normalized expression from 21–24 transcriptomes per condition (8 conditions) used to generate global transcriptome profiles. Transcripts Per Million (TPM) sequenced used to calculate mean expression. [b] PCA separates transcriptomes between invasive and native genotypes. Three genotypes per subspecies and each genotype is represented by 6-8 biological replicates. Transcriptomes from rhizomes cluster more distinctly between cut and uncut samples than those in shoots. [c] Pearson’s correlation of basal expression between invasive and native 1:1 orthologs. Colored regions highlight biased expression ≥2 SD in 31,854 ortholog pairs. Selected genes that are highly expressed in each group is marked. [d] Representative functional classes of genes with high and biased expression (marked by colored regions in [c]) in invasive and native tissues. Orthologs were classified as highly expressed and biased if mean normalized expression was Log2 ≥ 5 in one subspecies while ≤ 3 in the other.

#### High basal expression

Multiple genes associated with stress tolerance, energy metabolism, and growth regulation had higher basal expression in the invasive transcriptomes than in the native (Fig. 2c). Notably, *NADH dehydrogenase B1 (NDB1), δ1-pyrroline-5-carboxylate synthetase (P5CS2),* and *Auxin-resistance protein (AXR1)* were highly expressed in rhizomes both before and after cutback, as well as in shoots, but only in invasive genotypes. The corresponding orthologs in native genotypes remained minimally expressed (Fig. 2c). *NDB1* aids in stress tolerance by maintaining mitochondrial redox balance, reducing ROS accumulation, and providing flexibility in respiratory metabolism (Rasmusson et al., 2004). *P5CS* is the rate-limiting enzyme of proline biosynthesis, central to plant responses to abiotic stresses such as ROS, drought, and salinity, and plays an essential role in growth and development (Szabados and Savouré, 2010). *AXR1* mediates auxin signaling and promotes cytokinin responses, with broad pleiotropic effects on growth and development (Li et al., 2013). Additional stress-associated genes, including *CBL-interacting protein kinase (CIPK16)* (Tripathi et al., 2009), *Late embryogenesis abundant protein (LEA3)* (Liang et al., 2019), *adenosine 5’-phosphosulfate reductase-like protein 3 (APRL3)* (Kopriva and Koprivova, 2004), and *protein disulfide-isomerase 1 (PDIL1)* (Xia et al., 2018) were also highly expressed specifically in invasive rhizomes prior to cutback.

Elevated basal transcript levels in invasive rhizomes indicate a “stress-ready” state, where heightened energy metabolism and stress-protective pathways may enhance resilience following cutback. This is evidenced by functional biases among a few genes highly expressed in invasive, but not native, uncut rhizomes. We further tested whether this bias extends to all 1:1 orthologs with divergent expression between subspecies (Table S3.4 defined as those ≥2 SD from the Pearson correlation (Fig. 2c). In invasive tissues, especially uncut rhizomes, genes with higher basal expression were enriched in primary metabolism (amino acid, sugar, and fatty acid pathways), indicating increased availability of structural and energy precursors relative to native rhizomes (Fig. 2d) that can better sustain growth following damage.

### Subgenome-expression bias and retrotransposon-driven gene gain reinforce stress-resilient expression in invasive *P. australis*

The invasive and native genomes have retained unequal proportions of homeologs across their subgenomes and exhibit disrupted synteny between subgenomes (Figure 3a), with differential fractionation and transposable element activity further amplifying subspecies differences (Fig. 1b,c,d). These localized genomic changes likely contribute to expression divergence and, consequently, trait variation. To assess these differences, we examined lineage-specific expression associated with genes lacking direct orthologs in the other subspecies that included: (1) duplicated or fractionated genes with unequal homeolog retention in subgenomes, (2) novel genes arising from lineage-specific transposable element activity, and (3) genes present in the B chromosome in the invasive genome.

**Figure 3.**
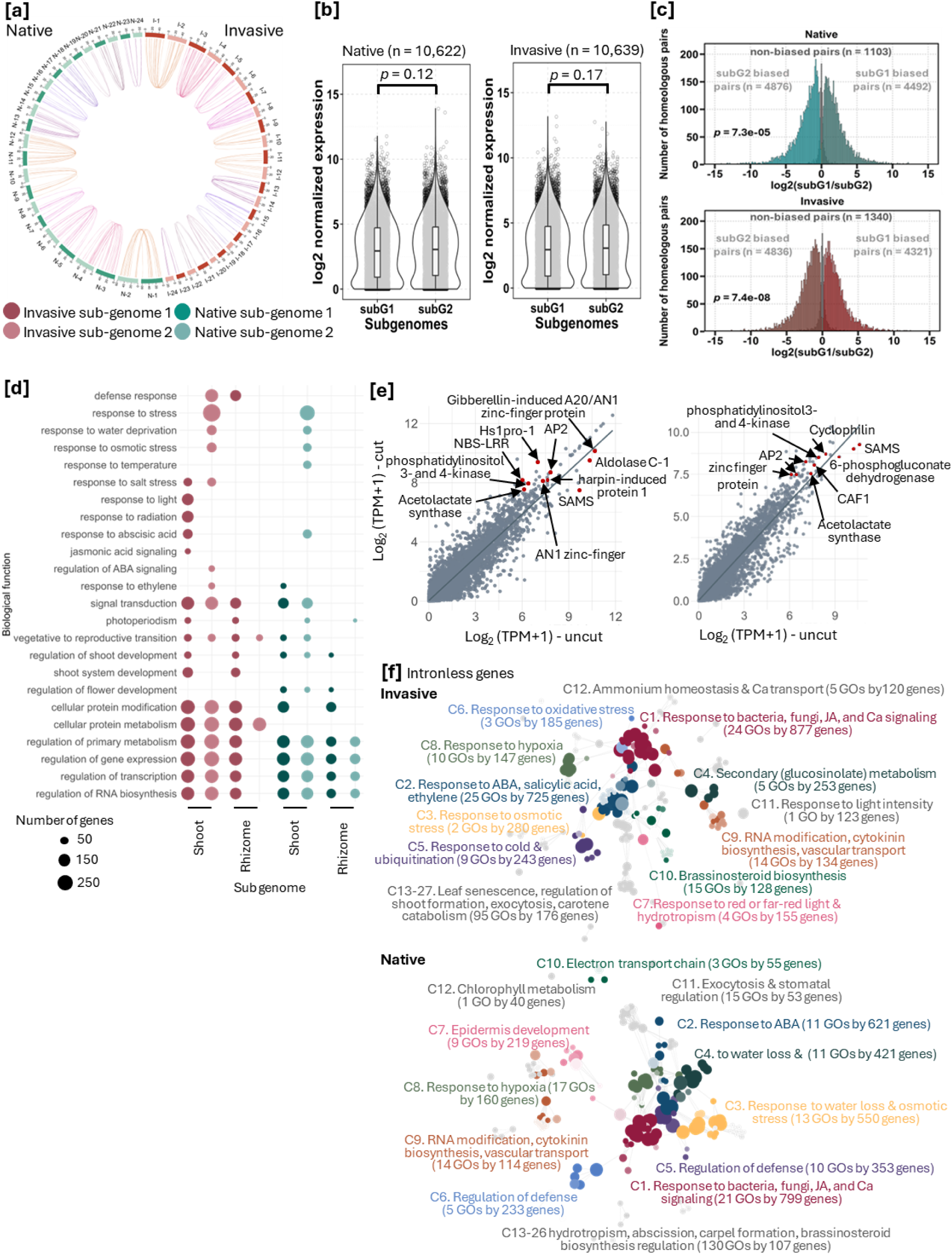
Subgenome bias and intronless gene expansion enhance stress responses in invasive *P. australis*. [a] Homeolog pairs between subgenomes 1 and 2 within a subspecies show preserved colinearity. Colored lines connect homeologs between subgenomes. [b] Global mean expression of homoeologs between subgenomes. Homeolog expression from 12 conditions representing 3 genotypes (invasive I1-3 and native N1-3) were pooled to get the global mean expression. p-values from paired Student’s t-tests, corrected using the Benjamini–Hochberg method. [c] Expression bias among homeolog pairs that had higher expression in subgenome1, in subgenome2, or unbiased expression in both subgenomes. Red regions indicate subgenome1 (subG1)-biased homoeologs with log_2_ expression fold change > 0 and blue regions indicate subgenome2 (subG2) with log_2_ expression fold change < 0 at Benjamini-Hochberg adjusted p-value < 0.01 based on paired Student t-test. Chi-square test was used to determine whether the observed number of biased homeologs significantly deviates from equal sub-genome representation, and the Benjamini-Hochberg adjusted p-values are shown for each comparison. [d] Enriched functions represented by homeologs dominant in subgeneome 1 and subgenome2 that are differently expressed in cutback compared to uncut samples. All displayed functional categories exclude overly broad (>2,000 genes) or very narrow (<10 genes) functions, while retaining non-redundant representative processes. All enriched functions have adj p-values <0.05. [e] Expression of intronless genes in invasive (8311 genes) and native (7651) rhizomes. Red data points highlight highly expressed genes known for stress responses. [f] Functionally enriched processes represented by intronless genes. Nodes represent GO terms, with size proportional to gene number; colors group similar functions and reflect significance (darker = lower adjusted p). Edges indicate shared ortholog families between functions. All clusters shown have FDR-adjusted p < 0.05.

#### Asymmetric subgenome homeolog expression

The two subgenomes show similar chromosomal compositions (Figure 3a) but within each subspecies, homeologs from the two subgenomes exhibit gene duplication or loss, interrupting the overall synteny (Fig. 1d). Based on higher retention of duplicated copies compared to rice, Wang *et al*. reported that subgenome 2 (also referred to as subgenome D) in *P. australis* (collected from Yuncheng, Shanxi province, China) was dominant in gene retention following fractionation (Wang et al., 2024b). Here, subgenome 1 in the invasive *P. australis* genome contained more genes than subgenome 2, but the opposite pattern occurred in the native genome (Table 1). Because fractionation is an ongoing process, and because pseudogenization cannot be reliably inferred from synteny with a distant diploid relative such as rice, we examined whether gene expression showed subgenome dominance and whether this contributes to adaptive traits in the invasive subspecies. The global mean expression of genes assigned to each subgenome did not differ significantly between subgenomes 1 and 2 in either subspecies (Fig. 3b) and was consistent when transcriptomes were pooled by genotype within each subspecies (Figure S5). However, when expression was analyzed across the 12 homeologous chromosome pairs, a greater number of genes showed higher expression from subgenome 2 than from its homeolog in subgenome 1 in all genotypes examined (Figure S6). This trend persisted when three genotypes per subspecies each were pooled confirming expression bias favoring subgenome 2 across both invasive and native *P. australis* genomes (Fig. 3c). We next tested whether homeolog pairs exhibiting subgenome-biased expression were associated with specific functional classes (Fig. 3d, Table S3.5). Expression biases for genes involved in primary metabolism, transcription, and translation were similarly distributed across subgenomes in both subspecies, but genes associated with biotic and abiotic stress responses showed a pronounced bias toward subgenome 2, particularly in invasive shoots. By contrast, genes associated with hormone responses were biased toward subgenome 1. These results suggest complementary functional specialization between the two subgenomes where subgenome 2 broadly reinforces stress-response pathways, whereas subgenome 1 is associated with hormone-mediated regulation, shoot development, and light responses (Fig. 3d).

#### Retrotransposon-mediated subspecies-biased gene expression

Retrotransposon-mediated reverse transcription of mature mRNA into DNA, followed by reintegration into the genome, is the most common mechanism generating intronless genes in grass genomes (Chen et al., 2023; Wang et al., 2006). To determine whether there is expression bias introduced by retrotransposon-driven gene insertions, we used intronless genes as proxies and compared their abundance and expression between invasive and native genomes. Retrotransposon-driven gene gains can occur in the ancestral lineage prior to subspecies divergence and independently within each lineage after their divergence from the tetraploid *P. australis* ancestor. Highly-expressed intronless genes in invasive *P. australis* exhibited higher expression levels than in the native genome and were more strongly induced in cut tissues compared to uncut controls (rhizomes (Fig. 3e), shoots (Figure S7a, b)). Functional annotations from homologs in model plants indicated that most highly expressed intronless genes were linked to biotic or abiotic stress responses including S-adenosylmethionine synthetase (SAMS), NBS-LRR proteins, and cyclophilins (Fig. 3e, S7a,b). To evaluate whether stress-related functions were broadly enriched among intronless genes, we tested for functional enrichment in both subspecies. In invasive *P. australis*, functionally-annotated intronless genes were enriched for abiotic and biotic defense pathways and hormone signaling networks regulating a wide range of stress responses (Fig. 3f). 1,729 invasive intronless genes fell into stress– or defense-related categories compared to 1,130 associated with growth and metabolism (Fig. 3f). By contrast, the native had fewer intronless genes linked to stress-responsive functions (1,688) or growth-promoting functions (290) (Fig. 3f). In both subspecies, stress-associated intronless genes showed higher mean expression than those involved in general metabolism in uncut shoots and in cut rhizomes (Fig. S7c, d). Together, these results demonstrate that highly-expressed intronless genes have preferentially reinforced stress-responses and growth-promoting pathways in the invasive subspecies.

#### Invasive-specific expression of B chromosome genes

B chromosomes are relatively small, non-essential chromosomes typically enriched in repetitive elements with few protein-coding genes primarily associated with chromosome segregation, as reported in several *P. australis* ecotypes, maize, and rye (Blavet et al., 2021; Chen et al., 2024; Cui et al., 2025; Wang et al., 2024b). Consistent with these findings, here the B chromosome in invasive *P. australis* subspecies is largely composed of repetitive sequences and uncharacterized proteins (Fig. S3, File S5). We found that 95 B chromosome genes were expressed in the sampled tissues (Fig. S8b, Table S5.1, comparable to 88 expressed genes reported for the maize B chromosome (Blavet et al., 2021). We then assessed functional biases among coding genes on the B chromosome and identified two major categories: photosynthesis and chromosome segregation during meiosis (Figure S8a. Table S5.1). However, these expressed genes are unlikely to contribute directly to invasive traits but may reflect mechanisms that promote B chromosome maintenance across generations.

### Transcriptional reprogramming prioritizes growth and stress tolerance to drive rapid regrowth in invasive *P. australis* after cutback

We expected significant transcriptomic shifts and structural and metabolic reprogramming needed for recovery after cutback. Shared responses between subspecies would reveal core processes for regrowth, while invasive-specific responses would highlight enhanced resilience to cutback and other physical damage. We identified differentially-expressed genes (DEGs) by comparing cut to uncut samples within each genotypes per subspecies for shoots and rhizomes, resulting in four transcriptome comparison groups for both invasive and native samples. A gene was considered a DEG for a subspecies only if it was independently identified as significantly different (padj < 0.05) in at least two of the three genotypes examined. We identified 20,109 (shoot, Table S6.1) and 14,963 (rhizome, Table S6.2) DEGs in invasive subspecies, compared to 16,499 (shoot, Table S6.3) and 12,034 (rhizome, Table S6.4) DEGs in native subspecies, indicating a stronger transcriptional response (∼23% increase) in the invasive for both tissues in response to cutback (Figure S9). Moreover, genes that were concordantly up– or down-regulated within a subspecies outnumbered those showing unique or discordant expression (Fig. S9).

#### Transcriptional alignment to promote growth

To assess transcriptional readjustment between growth and stress resilience, we categorized enriched functions among DEGs into broad classes representing cellular development, plant growth, hormone regulation, primary metabolism, and stress (Table S6.5, S6.6; Figure S10). Growth– and development-related genes were broadly downregulated following cutback, except in invasive rhizomes, which instead showed strong induction of genes associated with shoot meristem development (Figure 4a,b, Fig. S10). DEGs in invasive rhizomes shifted toward growth and hormone regulation, whereas native rhizomes showed a bias toward stress responses (Fig. 4a). DEGs associated with stress responses outnumbered those linked to growth in shoots of both subspecies (Fig. 4b). However, the invasive rhizomes uniquely upregulated genes associated with sulfur and nitrogen starvation responses, suggesting enhanced nutrient acquisition and ion transport, which was not observed in native rhizomes (Fig. S10a). Invasive shoots, unlike native shoots, showed greater induction of gibberellic acid–associated genes, a hormone that promotes internode elongation, leaf expansion, sustained meristem activity, and developmental transitions (Fig. 4b).

**Figure 4.**
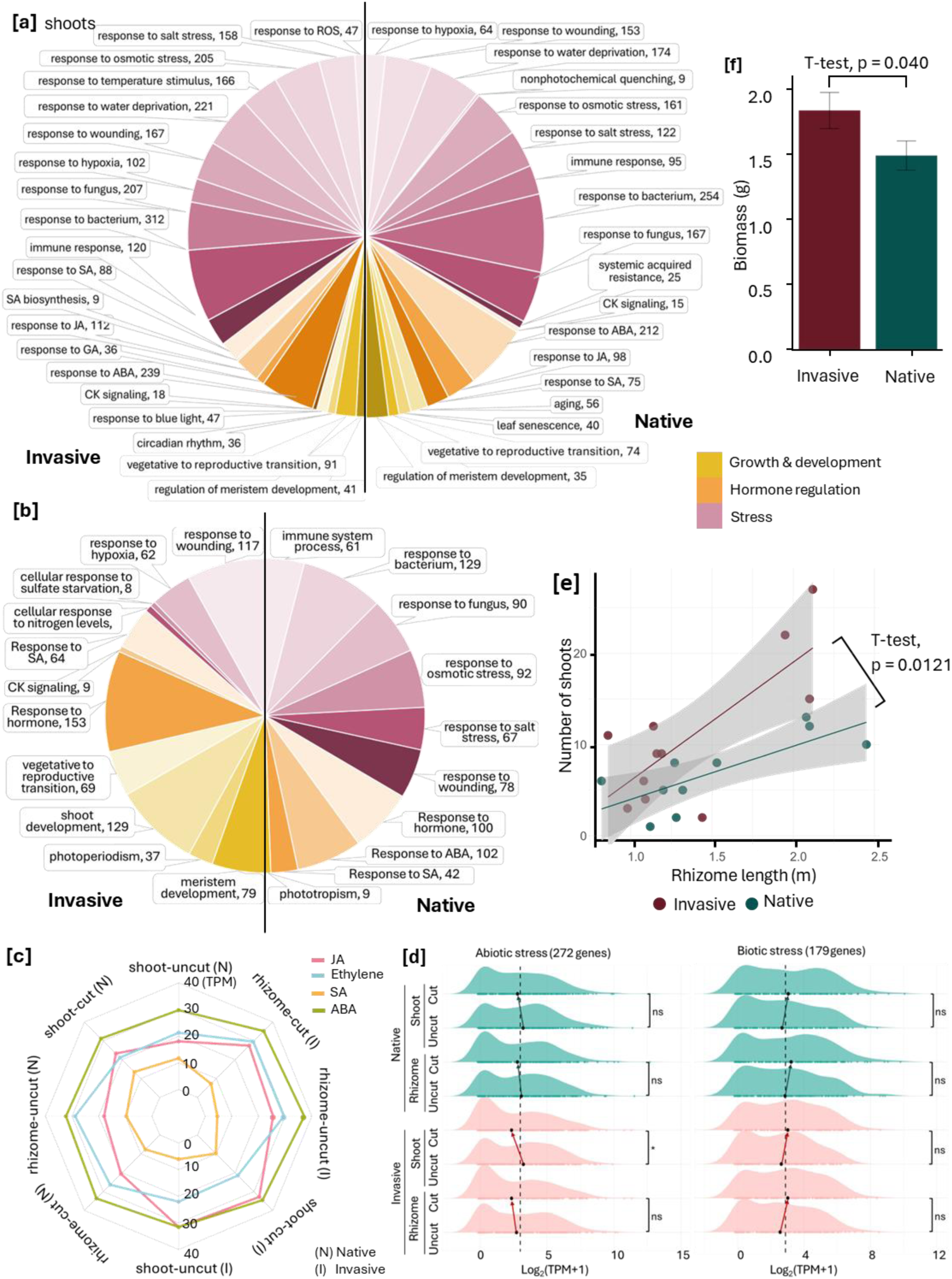
Cutback-induced regrowth in invasive rhizomes is supported by tissue-specific transcriptional reprogramming. Enriched functions associated with differentially expressed genes (DEGs) induced or suppressed after cutback in [a] shoots and [b] rhizomes. DEGs between pairwise comparisons of cut and uncut tissue were determined by p-adj values ≤ 0.05 based on Benjamini-Hochberg correction for multiple testing. Functional enrichment tested at adj p-values <0.05 with false discovery rate correction applied. The number of DEGs represented by a function included with each function. [c] Expression of *P. australis* homologs expressed at >10 TPM at least in one condition annotated under Salicylic Acid (SA) associated pathways (GO:0009751: Response to SA, GO:0009696: SA metabolism, GO:0009863: SA signaling), Jasmonic Acid (JA) associated pathways (GO:0009753: Response to JA, GO:0009695: JA biosynthesis, GO:0009867: JA signaling), Abscisic Acid (ABA) associated pathways (GO:0009737: Response to ABA, GO:0009688: ABA biosynthesis, GO:0009738: ABA signaling), Ethylene associated pathways (GO:0009723: Response to ethylene, GO:0009693: Ethylene biosynthesis, GO:0009873: Ethylene signaling). [d] Expression of P. australis homologs broadly annotated under Response to abiotic (GO:0006970: response to osmotic stress, GO:0006979: response to oxidative stress, GO:0009651: response to salt stress, GO:0009414: response to water deprivation, GO:0009408: response to heat, GO:0009409: response to cold, GO:0001666: response to hypoxia) and biotic (GO:0006955: immune response, GO:0009627: systemic acquired resistance, GO:0050832: defense response to fungus, GO:0042742: defense response to bacterium, GO:0051607: defense response to virus, GO:0002213: defense response to insect, GO:0002215, defense response to nematode) stresses. [e] Invasive rhizomes grow faster and produce more shoots than native rhizomes following cutback. [f] Biomass gain after another subsequent cutback is higher for the invasive subspecies compared to the native. Field-harvested rhizomes from native and invasive populations were cut into comparable segments and grown in pots for 48 days in growth chambers to assess growth.

#### Balance between abiotic and biotic stress responses

Invasive shoots exhibited a higher number of DEGs associated with abiotic stress responses, including wounding, reactive oxygen species, osmotic, and salt stress. They also showed greater induction of pathogen defense genes (Fig. 4a,b). Physical damage to cell membranes can cause the rapid release of stored fatty acids, particularly linolenic acid, which are used for oxylipin biosynthesis. Jasmonic acid (JA) is the most prominent oxylipin metabolite and functions as a potent intracellular signal that rapidly accumulates and spreads systemically to activate broad defense responses against both abiotic and biotic stresses (Hou et al., 2016; Lim et al., 2017). We observed strong transcriptomic signatures of DEGs associated with lipid catabolism and induction of JA-responsive defense genes in both invasive and native shoots following cutback (Fig. 4a,b, Fig. S10). However, invasive shoots had more upregulated genes tied to both biotic defense (e.g., immune responses to bacteria and fungi, systemic acquired resistance) and abiotic stress (e.g., salt, osmotic, water loss, hypoxia, and wounding) than native shoots (Fig. S10). These results suggest that invasive *P. australis* adopts a transcriptional reprogramming strategy that balances stress preparedness with sustained growth following cutback, supported by enhanced meristem activity, nutrient acquisition, and coordinated activation of biotic and abiotic defense pathways.

We next tested whether hormone-mediated transcriptional allocation between abiotic and biotic stress responses is a molecular phenotype observed at the global transcriptome level (Fig. 4c,d) or primarily driven by targeted induction and suppression of specific DEGs (Fig. 4a,b), and whether this differs between invasive and native subspecies following cutback. We first examined average expression levels of genes associated with four major stress-related plant hormones: abscisic acid (ABA), ethylene, JA, and salicylic acid (SA), including genes involved in their biosynthesis, signaling, and downstream responses. Genes associated with ABA were consistently expressed at higher levels in both tissues of the invasive subspecies (Fig. 4c). JA-associated genes were higher in both cut and uncut invasive shoots, whereas SA-associated genes were generally higher in cut and uncut tissues of the native subspecies, except in cut invasive shoots (Fig. 4c). Genes associated with ethylene responses are specifically highly expressed in invasive rhizomes relative to shoots, and to native rhizomes (Fig. 4c). Many ethylene-associated genes were also upregulated upon cutback in the invasive subspecies, under the functional group for response to hypoxia (Fig. S10). The overall proportional transcriptional allocation between abiotic and biotic defenses remained relatively stable between uncut and cut tissues in both subspecies. However, following cutback, expression of abiotic stress–related genes generally decreased while the transcriptional allocation to biotic defense genes increased, indicating a transcriptional readjustment to boost pathogen defense responses. This was observed across all pairwise comparisons, although it was statistically significant only in invasive shoots (Fig. 4d). Together, these results demonstrate that induction of biotic defense following cutback represents a consistent global transcriptional phenotype, whereas responses to abiotic stress are more prominently mediated through the targeted regulation of specific DEGs.

The tissue-specific transcriptional shift toward growth induction in rhizomes (Fig. 4a), even with a tradeoff for abiotic stress responses in shoots following cutback (Fig. 4d), may allow an effective resource reallocation needed for rapid regrowth in the invasive subspecies. To test this, we harvested rhizomes from the field at the same locations used for genome and transcriptome sampling (Fig. S1b) and grew them in a greenhouse to quantify shoot production per unit rhizome length. Consistent with the transcriptomic patterns, invasive rhizomes produced more shoots per unit rhizome length than native rhizomes (Fig. 4e). We then excised comparable rhizome segments without shoots (i.e., a second cutback) from each pot, measured biomass prior to cutback, and reweighed them after 48 days of regrowth. Consistent with transcriptomic results, invasive rhizomes accumulated significantly more biomass than native rhizomes (Fig. 4f).

### Global coexpression clusters highlight minimal root growth suppression while DEG-driven clusters suggest enhanced immune response and shoot growth induction in invasive *P. australis*

Most complex traits, such as stress tolerance and growth, are governed by coexpressed and coregulated gene networks within functional modules. While DEGs highlight strongly responding genes, consistent coexpression can also play key roles. Together, DEGs and their coexpressed partners can reveal functional clusters shaped by additive effects across gene networks. We aimed to identify key functional modules, comprising coordinately expressed, functionally related genes, including those not strongly differentially expressed, that are important for recovery from cutback. We analyzed coexpression networks in three steps. First, we constructed coexpression clusters using DEGs within each subspecies and examined enriched functions in clusters showing the strongest response to cutback, identifying modules primarily driven by DEGs. Second, we clustered all expressed genes in both subspecies to detect shared modules responding to cutback, independent of DEG status or orthology. Third, we clustered 1:1 ortholog pairs to identify coexpression clusters with diametric responses between subspecies, revealing genes inherited from a common *P. australis* ancestor that have diverged in expression direction.

#### DEG-driven coexpression clusters

We examined six invasive and four native coexpressed gene clusters in shoots and rhizomes of both subspecies (Figure 5a, Figure S11; Table S7.1). The first two clusters exhibited similar coexpression trends in both subspecies and showed the strongest responses to cutback (Fig. 5a). Cluster 1 genes declined in expression following cutback and were enriched for cell cycle and growth functions (Fig. 5a). This pattern reflects the removal of tissues containing dividing cells, leading to a drop in transcriptional investment in cell division, and indicating that plants had not returned to pre-cutback growth levels. Cluster 2 genes were more strongly induced following cutback in invasive tissues compared to native tissues. In the invasive subspecies, these genes were specifically enriched for immune defense responses and far-red light signaling, suggesting activation of shade-avoidance and other light-responsive growth pathways that promote rapid shoot elongation and recovery following cutback. Although a similar number of genes were coordinately induced in native tissues, they were not enriched for any annotated functions (Fig. 5a).

**Figure 5.**
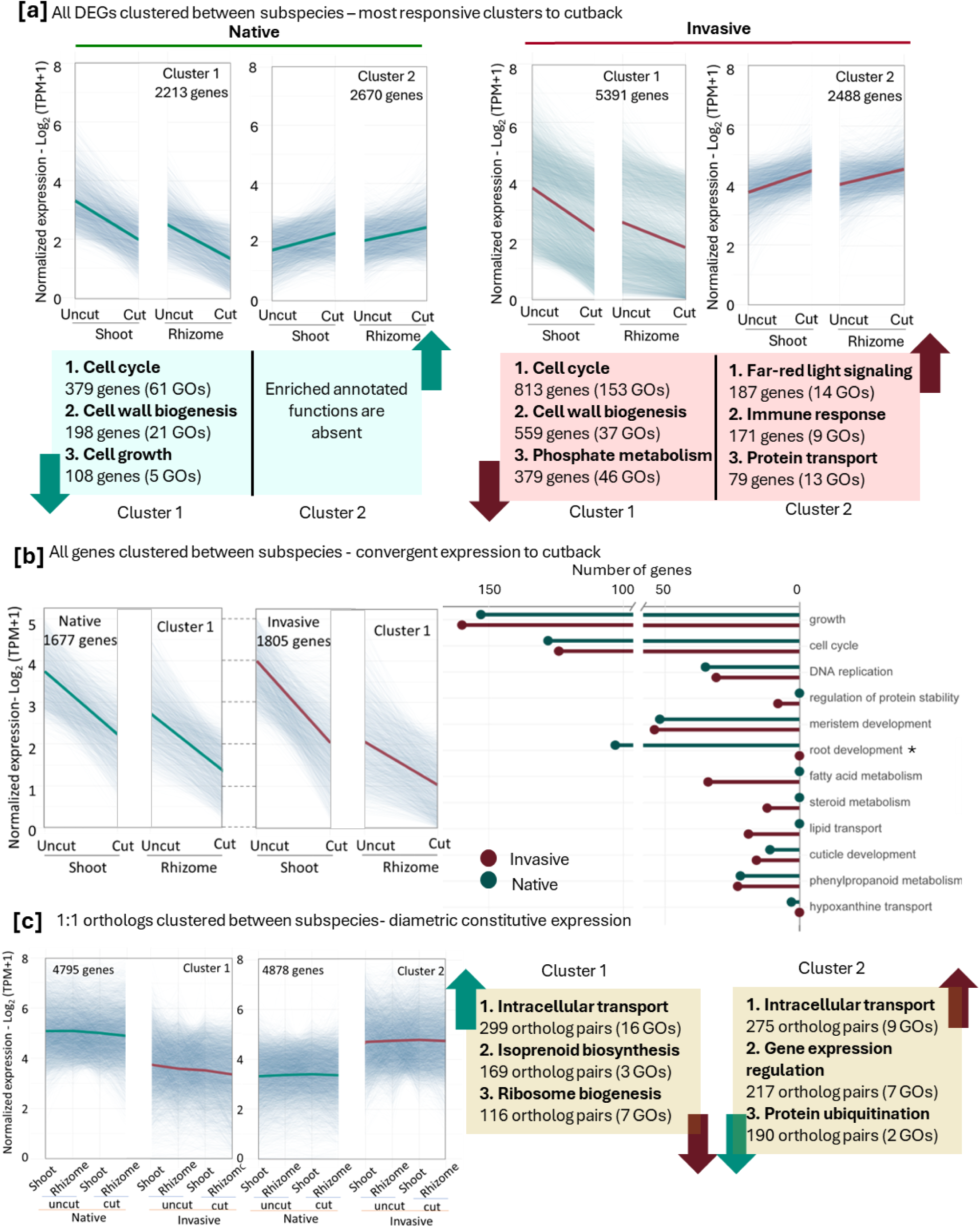
Functional insights from coexpression clusters reveal growth recovery and defense induction in invasive *P. australis*. [a] Each of the two gene coexpression clusters that showed the strongest response to cutback, identified independently using DEGs in native and invasive *P. australis*, along with the top three enriched representative functions for each cluster. [b] The coexpression cluster showing the strongest response to cutback in each subspecies, identified independently using all expressed genes and the enriched functions represented by the coexpressed genes in these two clusters. [c] Coexpressed clusters with diametric constitutive expression found with 1:1 orthologs between the subspecies and along with the top three enriched representative functions for each cluster. Coexpression clusters generated using k-means clustering performed with a membership cutoff of ≥ 0.5 across uncut and cut tissues from shoots and rhizomes. Thin lines represent individual genes contributing to the cluster while the thick line indicates the mean expression for the cluster. All functional categories exclude overly broad (>2,000 genes) or very narrow (<10 genes) functions, while retaining non-redundant representative processes as much as possible. All enriched functions have adj p-values <0.05 with false discovery rate correction applied. The top three functions represent those with the highest number of annotated genes, contributing most to the cluster.

#### Coexpression clusters shaped by all expressed genes

When all expressed genes were clustered across both subspecies, the strongest response to cutback was a sharp decline in expression in both subspecies (Fig. 5b, Figure S12; Table S7.2). Although Cluster 1 in each subspecies contained different numbers of genes, many functional categories were shared (Fig. 5b). In both subspecies, gene expression associated with cell cycle and growth were suppressed, further supporting an overall lag in growth after cutback, as suggested by DEG-driven clusters (Fig. 5a). However, the reduction in transcriptional investment in root development was markedly smaller in the invasive subspecies than in the native subspecies as reflected by the greater number of genes showing decline in the native subspecies (Fig. 5b). Interestingly, only ∼37% of Cluster 1 genes were 1:1 orthologs between the subspecies, indicating that lineage-specific transcriptional responses are at least as prominent as shared responses due to orthologous relationships, corroborating patterns observed in Figures 3 and 4.

#### Coexpression clusters of ortholog-pairs

Coexpressed 1:1 orthologs between the subspecies formed five clusters (Fig. 5c, Figure S13; Table S7.3). As expected for genes with conserved expression, three clusters showed consistent constitutive expression across tissues and subspecies (Fig. S13). In contrast, the first two clusters exhibited diametric expression patterns between the subspecies (Fig. 5c). Although these clusters were enriched for overlapping functions associated with cellular processes and not directly linked to traits biased towards an invasive lifestyle, their opposing expression patterns suggest divergence in regulatory controls specific to each subspecies.

### Evolutionary expansion of orthogroups highlights convergence for growth strategies and stress tolerance across invasive grasses

High resource allocation to root growth, sustained resource mobilization to growing sink tissues, optimization of light capture to maintain photosynthesis, and enhanced stress tolerance may provide a selective advantage for invasion success in *P. australis* (Fig. 1-5). We tested whether these genomic traits are consistently observed across invasive grasses relative to their non-invasive relatives (Figure 6), first by evaluating invasive *P. australis* compared to other grasses, by contrasting invasive and native *P. australis* relative to other grasses, and by assessing whether these traits are broadly prioritized across invasive grasses compared to non-invasive counterparts. We assessed support for evolutionary selection by analyzing orthogroup gene copy number variation relative to inferred ancestral gene family compositions at each node derived from extant genomes (Fig. 6a).

**Figure 6.**
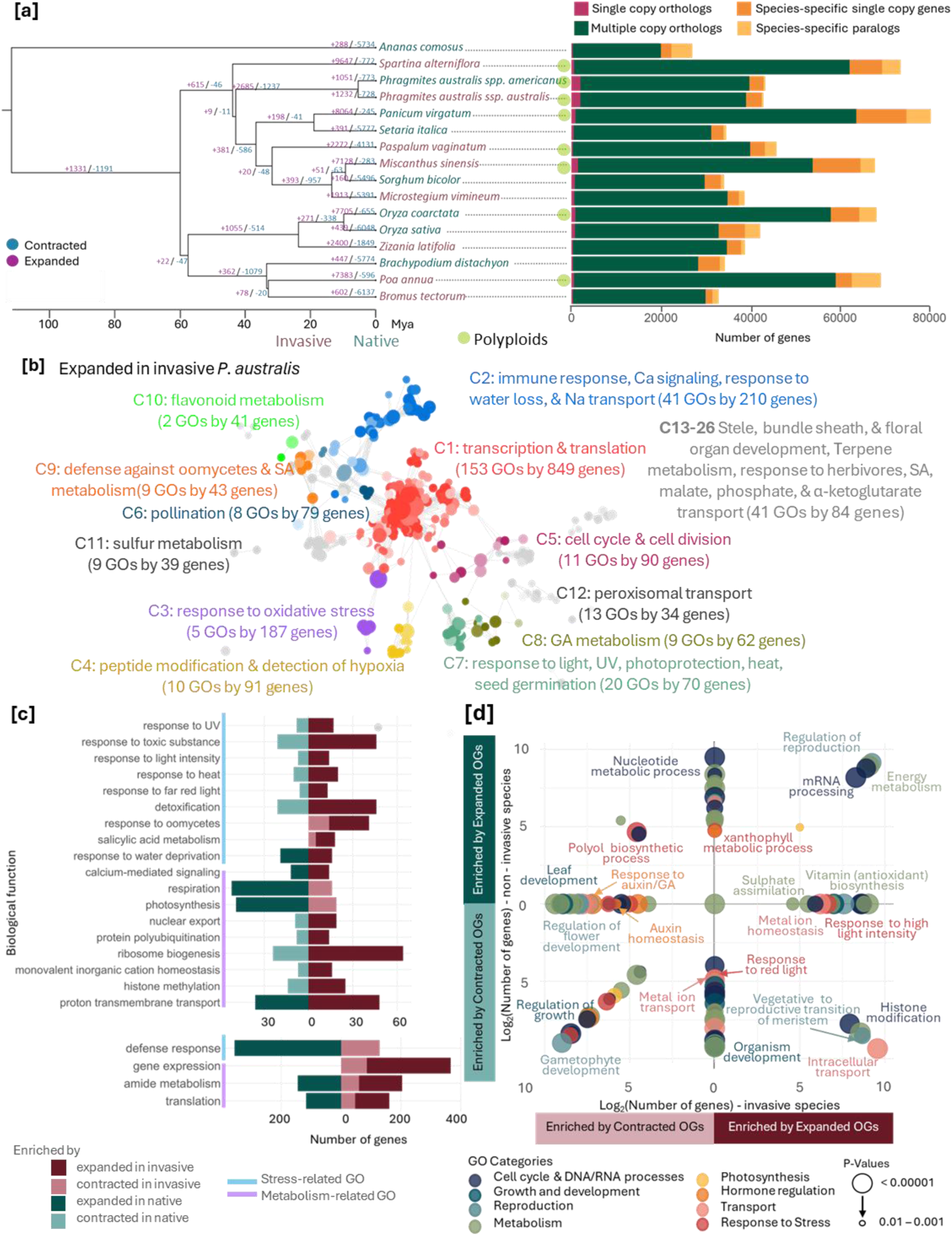
Growth and stress tolerance are shaped by orthogroup expansion and contraction in invasive versus non-invasive grasses. [a] Expanded and contracted orthogroups in 16 monocot genomes. Clade/linage relevant gene numbers are placed along branch lengths. The gene family composition for each genome is given next to each species. [b] Enriched functions of orthogroups expanded in the invasive *P. australis ssp. australis*. Nodes represent GO terms, with size proportional to gene number; colors group similar functions and reflect significance (darker = lower adjusted p). Edges indicate shared ortholog families between functions. All clusters shown have FDR-adjusted p < 0.05. [c] Shared functions between two or more groups associated with orthogroups that have expanded or contracted in invasive and native *P. australis*. [d] Enriched functions associated with gene families that are expanded or contracted in eight invasive grass species compared to their non-invasive relatives. Expanded and contracted orthogroups were counted from the divergent point relevant to each invasive-native pair. When the same gene function appeared in both expanded and contracted groups, net copy number variation was plotted. Biological processes were grouped into ten categories, each represented by a distinct color.

We used *P. australis* as the focal invasive–non-invasive pair and included seven additional grass species reported as invasive together with their closest non-invasive relatives (Fig. 6a, Table S8.1). These genomes were selected based on the completeness (presence of BUSCO orthologs >80%) and contiguity (i.e. available as chromosome-level genome assemblies) to ensure robust analyses of orthologous gene families (Figure S14). In total, our dataset was composed of 16 genomes, with a 50% representation from invasive species and *Ananas comosus* (pineapple) serving as the outgroup (Fig. 6a). Following the ancient rho whole-genome duplication shared by all Poaceae, lineage-specific WGD events have occurred frequently across grasses, including a polyploid origin reported ∼30 million years ago for the *P. australis* lineage (Oh et al., 2022; Wang et al., 2024b; Zhang et al., 2024). To capture a representative sample of grass genome diversity and to distinguish genomic traits facilitating invasiveness beyond signals associated with polyploidy, we included approximately equal proportions of polyploid (8) and diploid (7) species in our comparative analysis (Fig. S14). Multi-copy orthologs comprised the largest gene category across the genomes examined. Although polyploid genomes contained more protein-coding genes than diploids, neither higher number of genes nor polyploidy were biasedly associated with invasiveness (Fig. 6a). We identified orthologous gene families and assessed lineage-specific expansions and contractions (Fig. 6a and Table S8.2, S8.3).

#### Expanded orthogroups in invasive P. australis

The enriched functions of expanded gene families reflect a balanced investment in growth and stress tolerance (Fig. 6b). Selection towards retaining extra gene copies in families associated with light response, photoprotection, and bundle sheath development, along with malate, phosphate, and α-ketoglutarate transport and sulfur metabolism, suggest enhanced photosynthetic efficiency and nutrient flux to growing tissues. Peroxisomal transport and oxidative stress responses further maintain redox homeostasis during high photosynthetic and respiratory activity, preventing metabolic bottlenecks. Gene families associated with cell division and GA metabolism are also expanded, supporting coordinated growth with additional gene copies available for transcription and translation. Stress sensing and defense are similarly enhanced through expanded orthogroups contributing to Ca²⁺ signaling pathways, which mediate rapid responses to water deficit, Na⁺ stress, hypoxia, and oxidative cues, which are also associated with expanded families. SA-mediated defense and secondary metabolism, including flavonoid and terpene biosynthesis, and other pathogen– and herbivore-defense pathways, are concordantly expanded, reinforcing functional modules that integrate biotic and abiotic stress tolerance (Fig. 6b). These orthogroup expansions suggest a synergistic advantage in invasive *P. australis* enhancing antioxidant capacity, stabilizing cellular functions under stress, and enabling efficient resource allocation to growth and defense.

#### Orthogroups divergently expanded between invasive and native P. australis

Analysis of altered orthogroups between *P. australis* subspecies identifies genes selectively retained or lost since their divergence from a common ancestor (∼5 mya) (Fig. 6a). To examine divergent patterns, we analyzed reciprocal expansions and contractions of orthogroup sizes (Fig. 6b, Fig. S15) and identified functions shared across two or more of four categories (Fig. 6c, Table S8.4). We observed reciprocal patterns in orthogroups for light intensity, UV, heat stress, and ribosome biogenesis expanded in the invasive lineage but contracted in the native (Fig. 6c). Furthermore, transcription, SA metabolism, and response to oomycetes expanded in the invasive subspecies without a corresponding change in the native. Conversely, broad defense responses, particularly immune functions against multiple pathogen classes, were enriched among expanded families in the native but were reciprocally associated with contracted orthogroups in the invasive lineage (Fig. 6c). These results suggest that invasion success in *P. australis* may benefit from gene copy expansions that enhance performance under fluctuating light and abiotic stress rather than maintaining broad-spectrum defense gene families.

#### Recurrently altered orthogroups in invasive grasses

Compared to the *P. australis* focal pair, the other invasive species in our comparative analyses diverged from their non-invasive relatives much earlier and independently, spanning approximately 60 million years (Fig 6a). We tested whether any functional processes were predominantly associated with expanded gene families across eight invasive species, which would highlight convergent selection for invasive lifestyles. Only orthogroups present in at least 80% of target species (invasive or non-invasive) and those that were annotated in *P. australis* genomes were included in our analysis. Expanded and contracted orthogroups were assigned to each species relative to their divergence from the closest comparative lineage. In total, we found 7,567 expanded orthogroups (15,384 genes) and 3,274 contracted orthogroups (6,586 genes) across the eight invasive species. The eight non-invasive comparators showed 4,720 expanded (7,440 genes) and 6,998 contracted orthogroups (16,734 genes) (Table 8.3). Invasive species generally exhibited more gene family expansions (60% more) and fewer contractions (50% less) than non-invasive species, which showed the opposite trend. Responses to high light intensity, antioxidant biosynthesis, and metal ion homeostasis were enriched among orthogroups preferentially expanded in invasive species, without corresponding changes in the non-invasive group (Fig. 6d, Table S8.5). Notably, intracellular transport (e.g., multiple cation efflux family proteins and vacuolar ATP synthase) and vegetative to reproductive transitions were associated with expanded gene families in invasive species, while those functions were enriched among contracted orthogroups in the non-invasive group (Fig. 6d). Together, these findings indicate that invasive grasses frequently expanded gene families underlying abiotic stress resilience, light intensity response, and developmental transitions, while their non-invasive relatives showed no change or contraction in ortholog copy numbers associated with those functions (Fig. 6d).

## Discussion

Understanding how invasive plants achieve ecological dominance is essential for predicting and managing biological invasions. The invasive *P. australis* genome exhibits three distinct features: 1) single-copy orthologs (Fig. 1, 2) intronless genes characteristic of retrotransposon-mediated gene innovation, and 3) subgenome expression asymmetry (Fig. 3) to fuel stress tolerance and resilient growth. It also exhibits a stress-ready basal transcriptomic state, with higher constitutive expression of stress-, energy-, and growth-related genes than the native subspecies (Fig. 2). Following cutback, the invasive subspecies undergoes stronger transcriptional reprogramming and faster regrowth, reflected in a greater number of DEGs, increased shoot production, and higher biomass accumulation (Fig. 4). Coexpression analyses further identified coordinated gene modules as molecular phenotypes underlying complex traits, whose additive effects collectively support resource reallocation toward regrowth while reinforcing stress resilience during recovery from cutback (Fig. 5). Finally, we identified expanded orthogroups with elevated copy numbers in invasive *P. australis*, including those shared across a wide range of invasive grasses, suggesting convergent genomic strategies that may confer a selective advantage (Fig. 6). Together, these findings describe generalizable genomic mechanisms by which grasses evolve resilience and invasiveness.

Gene fractionation, rearrangement, and duplication accumulated over ∼30 million years within the ancestral allotetraploid lineage of *P. australis* have been further shaped by divergence between native and invasive subspecies over the past ∼5 million years (Cui et al., 2025; Oh et al., 2022; Wang et al., 2024b). In the invasive subspecies, single-copy orthologs are enriched for stress tolerance functions (Fig. 1) while intronless genes show a biased selection towards environmental stress responses, consistent with evidence that bursts of transposable element activity often coincide with stress exposure in plant evolution (Belyayev, 2014). For example, the heat-activated ONSEN retrotransposon in *Arabidopsis* is induced under elevated temperatures (Roquis et al., 2021), and LTR retrotransposons in barley (BARE-1) proliferate in response to water stress (Kalendar et al., 2000). Thus, stress-induced retrotransposon activity can act as a source of novel genetic variation subject to selection, and colonization of new habitats is expected to impose strong selective pressures. Consistent with this, invasive *P. australis* retained intronless genes, characteristic of retrotransposon origins (Chen et al., 2023), that are enriched for stress responses (Fig. 3e,f) and are induced following cutback. This characteristic suggests a selective advantage through enhanced capacity to respond to and tolerate diverse stress conditions.

The genomic analyses and results reported here are based on a single reference genome for each of the two subspecies, and their transcriptomes are from a limited number of population samples from the same region. We recognize that the limited number of genome and transcriptome samples restrict the ability to generalize across the continental distribution of each subspecies and definitively identify the mechanisms of invasiveness in *Phragmites australis* and other species. Nevertheless, the consistency of results and their general agreement with other invasive/native grass pairs support the idea that invasive genomes exhibit a consistent suite of genetic and physiological traits enabling their invasiveness. Future research should expand the sampling of native and invasive lineages across a wider geographical range to explore the genomic diversity within each subspecies towards generating a pangenome and confirm that our results represent consistent differences between native and invasive lineages. Our results provide a necessary baseline for future research investigating fundamental mechanisms of invasiveness.

Baker and Stebbins (1965) proposed that invasive species benefit by being general-purpose genotypes characterized by plasticity, robustness, and efficient resource use, enabling success across novel habitats (Baker and Stebbins, 1965; Barrett, 2015). Our results extend this framework by showing how genomic plasticity and novelty are reflected in the expansion of select gene families, divergence of homeologs and orthologs between subgenomes and subspecies, and enrichment of intronless genes alongside canonical ortholog gene families. These features are more strongly enriched in the invasive subspecies than the native subspecies for functions supporting survival and growth under fluctuating light, water, nutrient, and salinity conditions. This correlates with enhanced environmental tolerance and phenotypic plasticity achieved through the capacity to adjust growth, physiology, resource acquisition, and development in response to environmental change. Our genome-based findings also mirror a meta-analysis of 75 invasive–native species pairs, which found that invasive species exhibit greater phenotypic plasticity in physiological traits (Davidson et al., 2011). Elevated basal expression associated with “stress preparedness” is also more pronounced in the invasive subspecies, further supporting a genomically-encoded robustness to harvest light and sequester nutrients across diverse and stressful environments. Following cutback, transcriptional reprogramming reflects rapid adjustments to sustain root and shoot regrowth, particularly in invasive *P. australis*, although both subspecies experience an initial transcriptional setback. The induction of stress-tolerance genes that are significantly differentially expressed after cutback is further reinforced by coordinated gene families mediating both abiotic and biotic responses. Notably, in invasive *P. australis* expanded gene families contribute more strongly to abiotic stress tolerance, while biotic stress responses are maintained through basal or inducible transcriptional programs. These genomic features are consistent with invasive *P. australis* functioning as a general-purpose genotype.

Being a generalist and maintaining a genetic toolkit to grow under a wide array of stresses necessarily involves tradeoffs. Hormonal crosstalk plays a major role in regulating these growth–defense balances (He et al., 2022), and, as expected, much of the transcriptional reprogramming following cutback in *P. australis* is allocated to stress hormones (Fig. 4). Plants frequently exhibit tradeoffs in resource allocation, balancing growth and stress tolerance, as well as constitutive versus inducible responses (Kempel et al., 2011; Lundgren and Des Marais, 2020). *P. australis* appears to strike a balance between three major functional gene classes competing for resource allocation: growth, biotic stress tolerance, and abiotic stress responses. This balance is achieved in part through genome-level organization, where gene family expansions are biased toward abiotic stress-related functions in invasive *P. australis*, while gene families associated with biotic stress responses are comparatively contracted relative to the native subspecies (Fig. 6). This tradeoff is mitigated at the transcriptome level by elevated basal expression of genes involved in both biotic and abiotic stress responses (Fig. 1–3), reflecting a constitutive “stress-prepared” state. Following cutback, a three-way tradeoff is observed in the invasive lineage where transcriptional resources are diverted from abiotic stress responses toward rapid regrowth, while maintaining or increasing the expression associated with biotic defenses (Fig. 4). Most abiotic stress-response categories are downregulated in the invasive subspecies, whereas pathogen-defense genes are preferentially induced after cutback compared to the native subspecies. We observed this trend both at the level of differentially expressed as well as coexpressed functional modules. Growth is prioritized in the invasive subspecies through both constitutive and induced transcriptional activation of energy metabolism, GA-mediated signaling, light-responsive pathways, root development, and nutrient uptake (Fig. 4, 5, S10). Overall, this balance between growth and stress tolerance is achieved through a multi-level regulatory strategy combining (i) genome-level biases in gene family expansion and contraction, (ii) transcriptome-level modulation of high basal (constitutive) expression states, and (iii) inducible expression programs following cutback, enabling flexible allocation of resources among growth, abiotic stress, and defenses against biotic stresses.

Decades of control efforts targeting invasive *P. australis* indicate that herbicide application and mowing (i.e., cutback) are often ineffective in the long term (Bowe et al., 2024; Hazelton et al., 2014; Lindsay et al., 2023). As a panglobal generalist, *P. australis* is inherently predisposed to outcompete native flora in introduced habitats, particularly where climate change further compromises local ecosystem resilience (Eller et al., 2014). It is well known for its ability to withstand control treatments and recover partly due to large belowground reserves stored in rhizomes relative to native lineages (League et al., 2006). Both cutback and herbicide treatments can induce shoot regeneration. Such responses reflect inherent physiological tolerance and may pre-adapt populations for the evolution of herbicide resistance given that herbicide resistance in weeds has evolved rapidly and involves multi-copy genes (Gaines et al., 2020) and that many herbicides target core metabolic and regulatory pathways. On the one hand, most genes in *P. australis* are represented by multi-gene families (Fig. 6a) while on the other hand, herbicide resistance is increasingly associated with multi-gene families coding for key metabolic enzymes and those linked to detoxification and regulatory reprogramming. Therefore, it is challenging to develop control strategies for invasive *P. australis* that are both specific and sustainable while maintaining long-term effectiveness and minimal impact on the co-occurring native subspecies.

Another approach to invasive species control using RNAi-based gene silencing can target essential metabolic and photosynthetic genes in invasive plants, including *P. australis*, offering environmentally sustainable and potentially effective control strategies (Cintra et al., 2025; Ji et al., 2025). Successful implementation, however, requires high-quality genome resources to ensure target specificity. Because many essential functions are encoded by multi-gene families, RNAi designs should account for gene copy number and aim to silence multiple homologs within a family. Greater efficacy may be achieved by simultaneously targeting non-homologous genes that do not share sequence similarity but are strongly coexpressed to contribute to essential physiological and metabolic processes. The genomic resources generated in this study enable the design of such strategies by allowing precise identification of gene family composition, guiding multi-target RNAi design, and prioritizing co-expressed gene modules critical for plant performance (Table S2, 4, 6, 7, 8).

Our study reveals that invasion success in *P. australis* is supported by the integration of genomic architecture and transcriptional regulation, enabling simultaneous optimization of growth and stress tolerance. Expansion of gene families, coupled with constitutive and inducible expression programs, provides both robustness and flexibility in response to disturbance. These insights advance a genome-level understanding of plant invasiveness and highlight the need for control strategies that account for functional redundancy and regulatory plasticity. The genomic resources generated here provide a foundation for identifying and prioritizing molecular targets for the next generation of gene-based control measures.

## Materials and Methods

Figure S16 outlines the overview of the experimental design and Table 2 lists tools used for genome assembly, annotation, RNAseq analyses, and comparative genomic analysis with specific parameters used.

**Table 2.** Tools and programs used for Methods.

| Task | Tool/Program (version & parameters when changed from default) | Reference |
| --- | --- | --- |
| Raw sequence post-processing and filter for quality | SMRT Analysis (v2.3) | (Pacific Biosciences., n.d.) |
| Genome assembly- primary assembly of contigs | Flye (2.9) (estimated genome size at 1 GB) | (Kolmogorov et al., 2019) |
| Assembly polishing and contig improvement using short reads | Pilon | (Walker et al., 2014) |
| Use linkage information from TELL-Seq to build long-range scaffolding connections without alignments | LINKS | (Warren et al., 2015) |
| Scaffolding contigs with linked reads | ARCS | (Yeo et al., 2018) |
| Analyze loop-resolution and manual inspection of pseudomolecules to chromosome assignments | Juicer (v2), Juicebox (v. 1.11.08) | (Durand et al., 2016) |
| Initial Hi-C Scaffolding | 3D-DNA (v.180922) | (Dudchenko et al., 2017) |
| Correct errors of long reads to be used in gap closing | Racon (v.1.5) | (Vaser et al., 2017) |
| Close gaps in the scaffolded genome assembly using long reads | TGS-GapCloser (tgstype pb, ne) | (Xu et al., 2020) |
| Subgenome separation | Subphaser (v.1.2.6) (-c sg.config, -pre default, -k 13, -q 50) | (Jia et al., 2022) |
| Whole genome alignment to assess collinearity | D-GENIES | (Cabanettes and Klopp, 2018) |
| Alignment between genomes | Minimap2 | (Li, 2018) |
| Gene alignments to genome | mmseq (v.14.7e284) (max-seqs 50, -s 7.5, search-type 3) | (Steinegger and Söding, 2017) |
| De novo identification of genomic repeats | RepeatModeler | (Flynn et al., 2020) |
| Masks known genomic repeats | RepeatMasker | (Nishimura, 2000) |
| Combine RNAseq evidence from multiple samples to assist genome annotation | StringTie2 | (Kovaka et al., 2019) |
| Gene model annotation | MAKER2 | (Holt and Yandell, 2011) |
| Noncoding RNA annotation | cmscan of Infernal (v1.1.2)<br>(cut_ga, rfam , noali) | (Nawrocki and Eddy, 2013) |
| Genome alignments, gene, LTR density visualizations | circos v0.69-8 | (Krzywinski et al., 2009) |
| BUSCO assessments for genome completeness | BUSCO (V.5.2)<br>(lineage: embryophyta_odb10) | (Manni et al., 2021) |
| Gene alignments with the rice genome | mmseqs2 (v13.45111) (easy-search --search-type 2) | (Steinegger and Söding, 2017) |
| To compare *P. australis* genes to *Arabidopsis* proteins | mmseqs2 (option -s 7.5 for maximum sensitivity) | (Steinegger and Söding, 2017) |
| Gene annotation with rice genome | RAPDB (annotation database) IRGSP-1.0_2025-03-19 | (Kawahara et al., n.d.) |
| Functional enrichment tests | BiNGO plugging in Cytoscape (v.3.10.1) (pvalue - 0.05, OBO file from GO consortium, custom gene association file) | (Maere et al., 2005) |
| Clustering and summary of enriched functions | GOMCL (gosize 1000, Ct 0.5, I 1.5, hm, nw) | (Wang et al., 2020) |
| Quality checks and adapter/contaminant trimming for RNASeq data | FastP | (Chen et al., 2018) |
| Mapping RNASeq reads to the genome | HISAT2 (v2.2) | (Kim et al., 2019) |
| Reference guided transcript identification and counting | StringTie (v 2.1.5) | (Pertea et al., 2015) |
| Co-expression analysis | K mean clustering in R (v.4.3.2) |  |
| Normalize RNASeq data and identify differently expressed genes | DESeq2 | (Love et al., 2014) |
| Build orthogroups | Orthofinder (v.2.5.4) | (Emms and Kelly, 2019) |
| Gene family expansion and contraction estimates | CAFE (v.5) | (Mendes et al., 2021) |
| Tree visualization | FigTree (v.1.4.4) | (Rambaut, 2018) |
| Obtain Ortholog, paralog counts, and other custom counts | Custom python (3.9) and R (4.3.2) scripts |  |
| Plotting figures (e.g. PCA, correlation upset, coexpression, scatter, box, density plots, and statistics) | R (4.5.0) ggplot2, dplyr, base stats, multcompView, readr & rlang | (R Core Team, 2025) |

### Genome sequencing, assembly, and annotation

Genomic DNA was isolated from leaf tissue of a single greenhouse-propagated clone for each subspecies. Source material for the invasive subspecies was collected at 41.63120838° N, −83.2289166° W, and for the native subspecies at 41.614755° N, −83.197046° W within Ottawa National Wildlife Refuge, Ohio, USA. The invasive genotype was confirmed as haplotype M and the native genotype as haplotype E using PCR with haplotype-specific primers (Saltonstall, 2016). High-molecular-weight gDNA was extracted using a 2% CTAB protocol (Doyle and Doyle, 1987), sheared to ∼30 kb (g-TUBEs, Covaris), and used to generate SMRTbell libraries (PacBio). Libraries were sequenced on a Sequel IIe to yield 253 Gb (7.18 million reads; mean read length 20,558 bp) for the invasive subspecies and 224.5 Gb (8.56 million reads; mean read length 26,214 bp) for the native subspecies as CLR reads. The same source of DNA was used to prepare Hi-C libraries (Omni-C kit, Dovetail Genomics) and linked-read libraries (TELL-Seq, Universal Sequencing Technology). Libraries were pooled, quantified by qPCR, and sequenced on two SP lanes (150 bp paired-end) on an Illumina NovaSeq 6000. Hi-C sequencing generated 308 Gb (invasive) and 359 Gb (native) and TELL-Seq generated 253 Gb (invasive) and 307 Gb (native). The long-reads were used to generate the primary contig assembly which was further scaffolded using the linked and Hi-C reads. The assembled genome was masked for repeats and annotated for repeats and genes (Table 2).

### Plant material for comparative transcriptomics, RNA-seq, and physiological assessments

Rhizome cuttings were collected by the USGS Great Lakes Science Center from populations in Michigan and Ohio (Fig. S1b). The native subspecies *americanus* was easily distinguished from the invasive subspecies *australis* given its smaller stature and reddish stems with distinct floral and vegetative characteristics (Saltonstall et al., 2004). The native populations typically occur intermixed with more diverse plant communities whereas the invasive subspecies more typically formed large monocultures reflecting its great competitive ability (Saltonstall, 2003). Additionally, we reconfirmed that the native populations used in this study belonged to chloroplast haplotype E, while the invasive populations belonged to haplotype M, based on previously reported DNA markers (Lindsay et al., 2023). Cuttings were cleaned, resprouted in water for 4 weeks, and shoots ≥15 cm with roots were excised, leaving 3–5 cm of rhizome on either side of the node (first cut). These were planted in a 3:1:1 mixture of peat moss, sand, and potting soil and kept in a walk-in growth chamber (WE-1012 Percival) set at 21°C and 70% relative humidity with a 14:10 hour light:dark cycle for two months. Next, plants were divided into cut and uncut treatments. Cut samples were then trimmed to soil level (second cut) and allowed to regrow for two additional months, while uncut controls remained undisturbed. At harvest, shoots and rhizomes were collected for RNA-seq. Cut samples contained regenerated shoots, whereas uncut samples retained shoots produced after the first cut. Each treatment included three genotypes per subspecies with 6–8 biological replicates (Table S4.1). RNA extraction, library preparation, sequencing, and differential expression analyses followed methods described in Oh et al. (2021). We designated genes as differentially expressed only if they were independently identified as significant (padj < 0.05) in at least two of the three genotypes examined, each represented by 6–8 biological replicates. Functional predictions based on transcriptomic responses were made only for specific expression profiles that converged across 21–24 independently generated transcriptomes per tissue or treatment group within each subspecies (Table S4.1), to minimize the influence of isolated population-level heterogeneity on overall subspecies-level characteristics. Biomass and shoot regeneration were measured in parallel experiments to evaluate transcriptome-based growth predictions.

### Comparative genomics analysis among invasive and non-invasive monocots

Sixteen monocot genomes, including 15 grass species and one non-invasive outgroup, were selected for comparative genomic analysis following quality assessment (Table S8.1). Invasive species and their non-invasive relatives were selected based on categorical assignments reported by the Global Invasive Species Database developed and managed by the International Union for Conservation of Nature (IUCN) (https://www.iucngisd.org/gisd/howto.php) and the database for Natural Resources Conservation Service managed by the United States Department of Agriculture (https://plants.sc.egov.usda.gov/) (Table S8.1). Species selection was guided by phylogenetic relationships in the NCBI Taxonomy Browser (https://www.ncbi.nlm.nih.gov/datasets/taxonomy/browser/). Orthologous gene families were identified from all protein-coding genes annotated for each genome, and gene family expansion and contraction were analyzed in a phylogenetic context. Custom scripts in R (R Core Team, 2025) and Python were used for data processing, integration, and visualization.

## Supplementary figure captions

**Figure S1.**
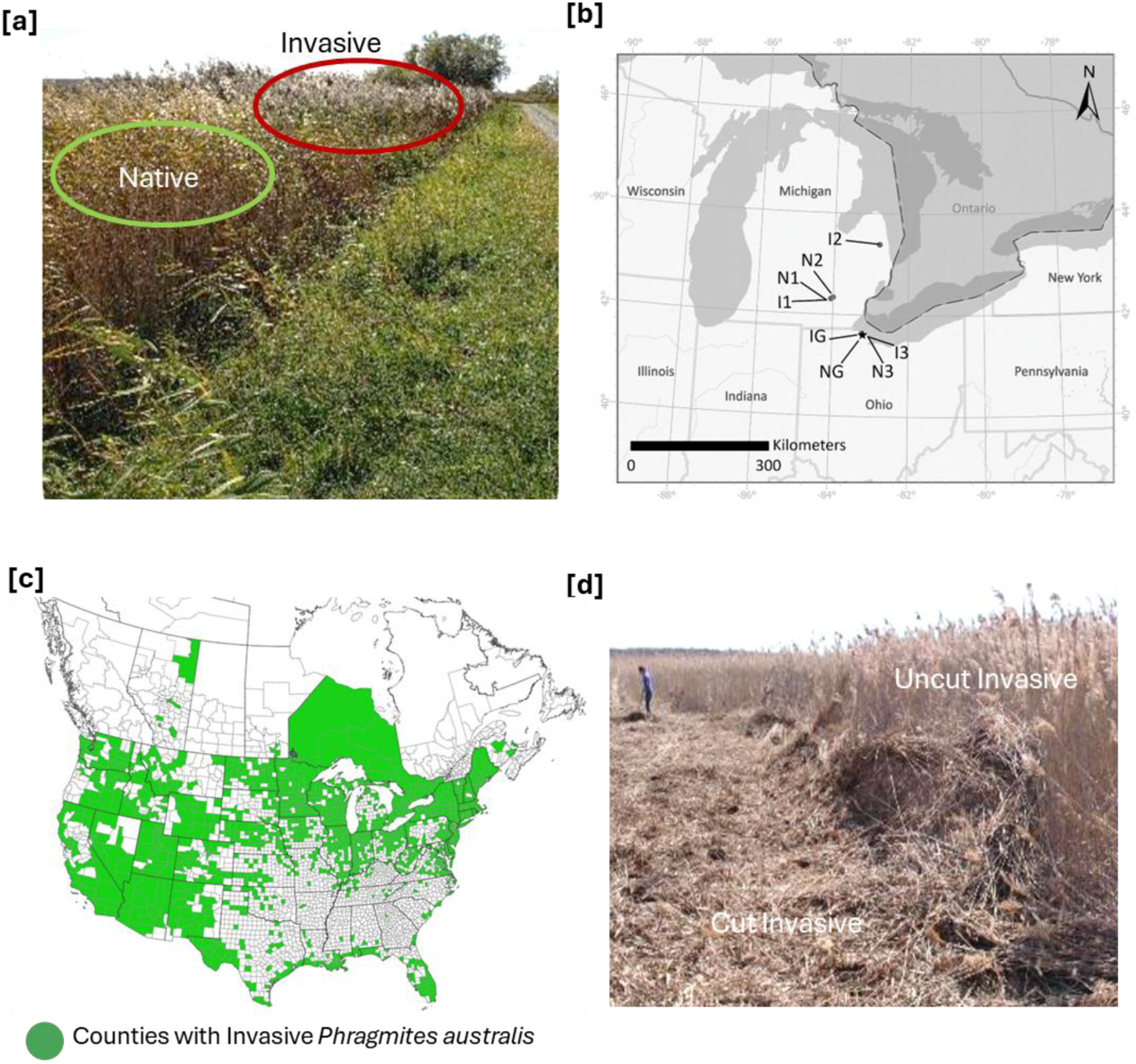
The invasive *Phragmites australis* populations, sampling sites, and a typical management strategy. [a] Tall invasive *Phragmites* (background) grows adjacent to a patch of native Phragmites (foreground) along a road at the U.S. Fish and Wildlife Service Ottawa National Wildlife Refuge, Oak Harbor, OH USA. Photographs by the U.S. Geological Survey. [b] Clonal rhizome cuttings for the reference genomes (IG and NG) were collected near the Ohio site. Sample collection sites, invasive (I1, I2, I3) and native (N1, N2, N3) for transcriptome analysis in Michigan (MI) and Ohio (OH) in the Great Lakes region. [c] Presence of invasive *P. australis* ssp. *australis* as reported by EDDMapS as of 6/24/2024. [d] Thick and dry aboveground biomass of invasive *Phragmites* is cut and piled to promote the growth of native plants at the Sandusky State Game Area, Sandusky, MI USA was cut similar to common management approaches.

**Figure S2.**
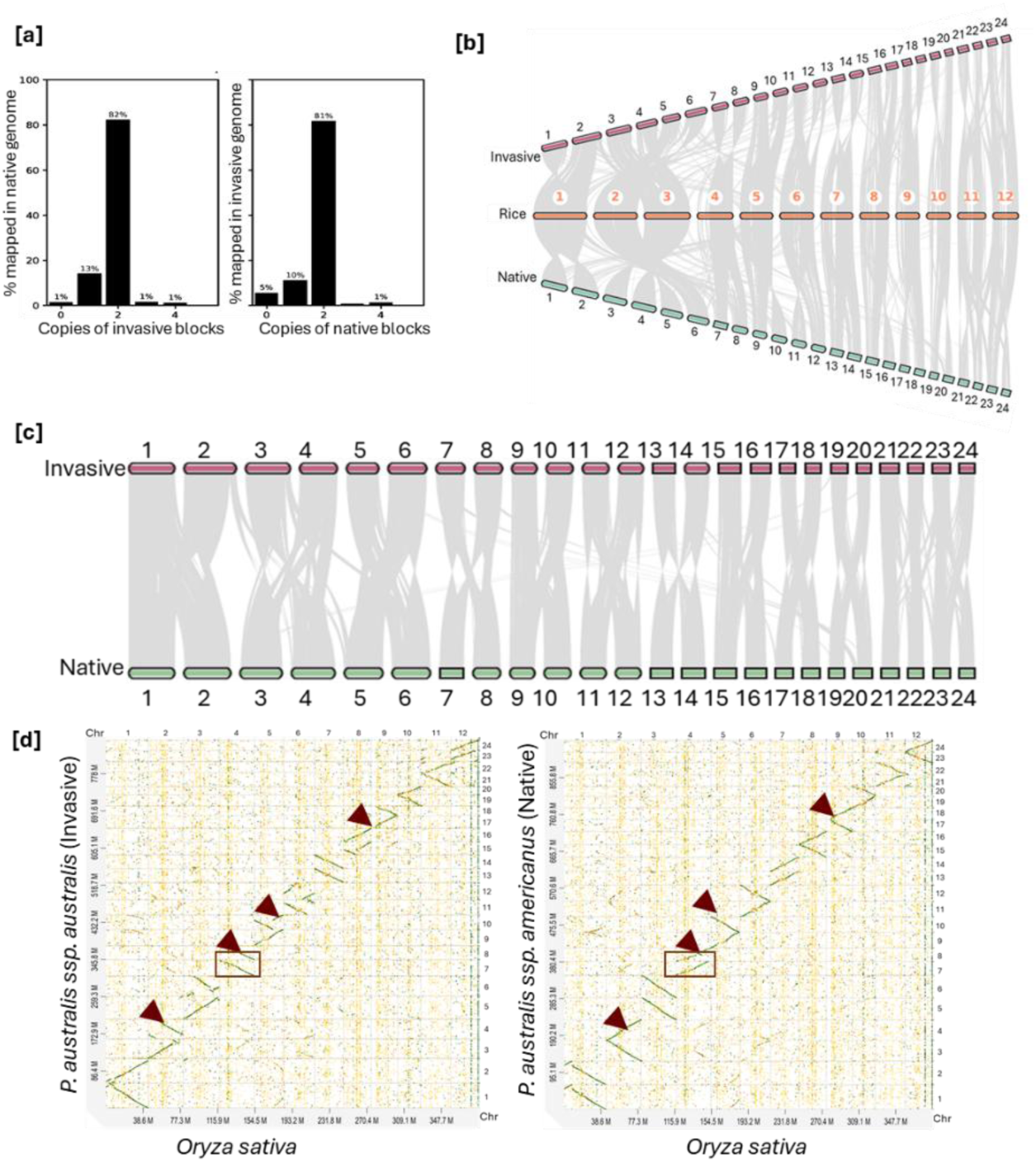
Subgenome organization of the tetraploid *P. australis* invasive and native reference genomes. [a] More than 80% of one genome shows duplicated genomic blocks in the other genome. [b] Twenty four chromosomes from invasive and native *P. australis* genomes aligned to the 12 chromosomes in the diploid rice genome (*Oryza sativa*). [c] The 12 chromosomes in each subgenome (even numbered set and the odd numbered set) aligns to the orthologous homeolog in the other genome. [d] Each genome aligned to the diploid Oryza sativa shows two *P. australis* chromosomes aligned to one chromosome *O. sativa* chromosome. Brown arrows mark examples of genome structural variation between invasive and native genomes when aligned to the *O. sativa* genome. The boxed region highlights shared similarity with *O. sativa* Chromosome 4, which appears as two duplicated regions colinear with Chromosomes 7 and 8 in the native *P. australis* genome, but as duplicated and inverted in the invasive *P. australis* genome.

**Figure S3.**
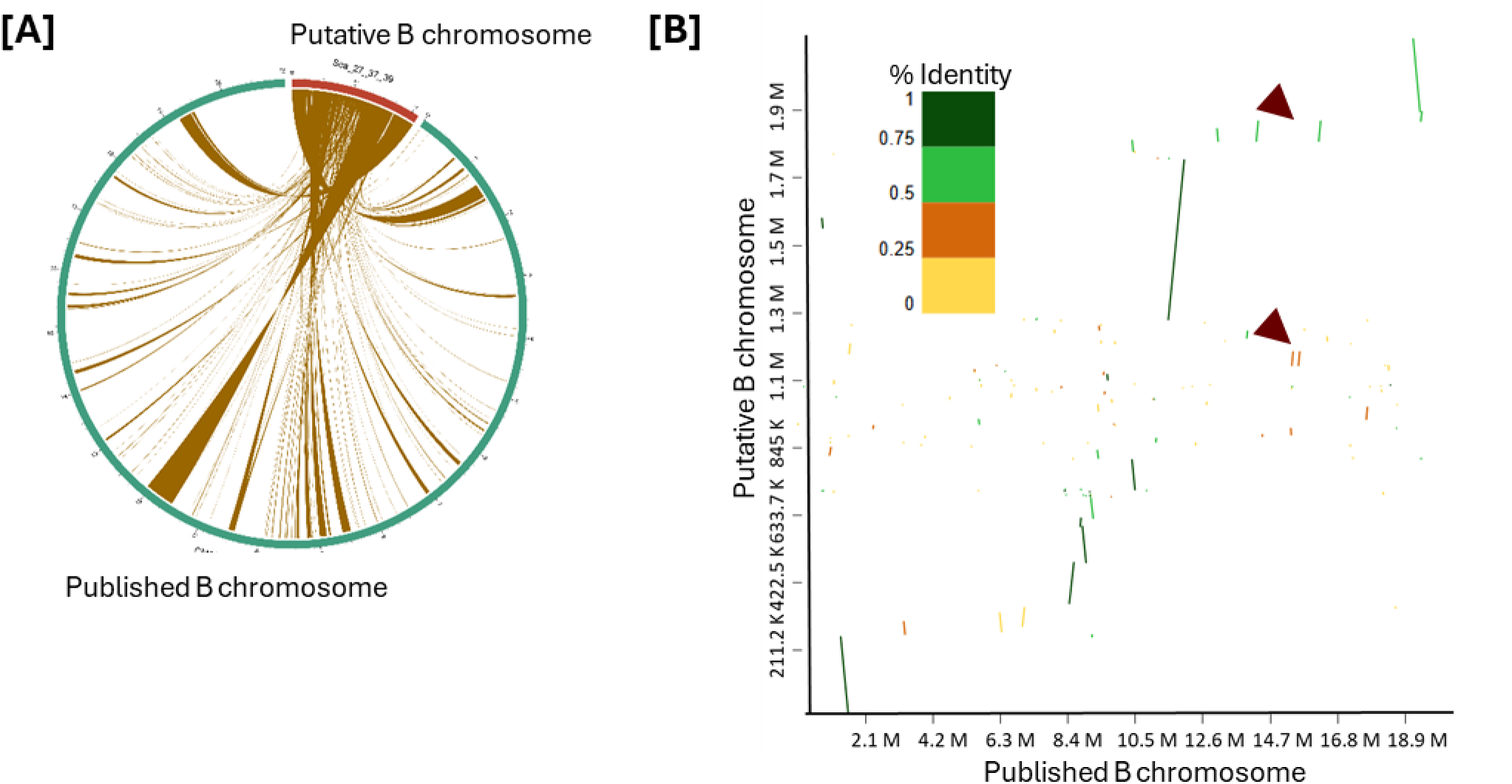
The putative B chromosome of invasive *P. australis* genome. [A] Contiguous alignment of the assembled B chromosome (magenta) to non-contiguous regions in the reference B chromosome reported by Wang, et al. (2024b) (green). [B] The alignment identity differs along the length of the two chromosomes, and some smaller regions are deleted or duplicated (examples shown by brown arrows) in one of the chromosomes.

**Figure S4.**
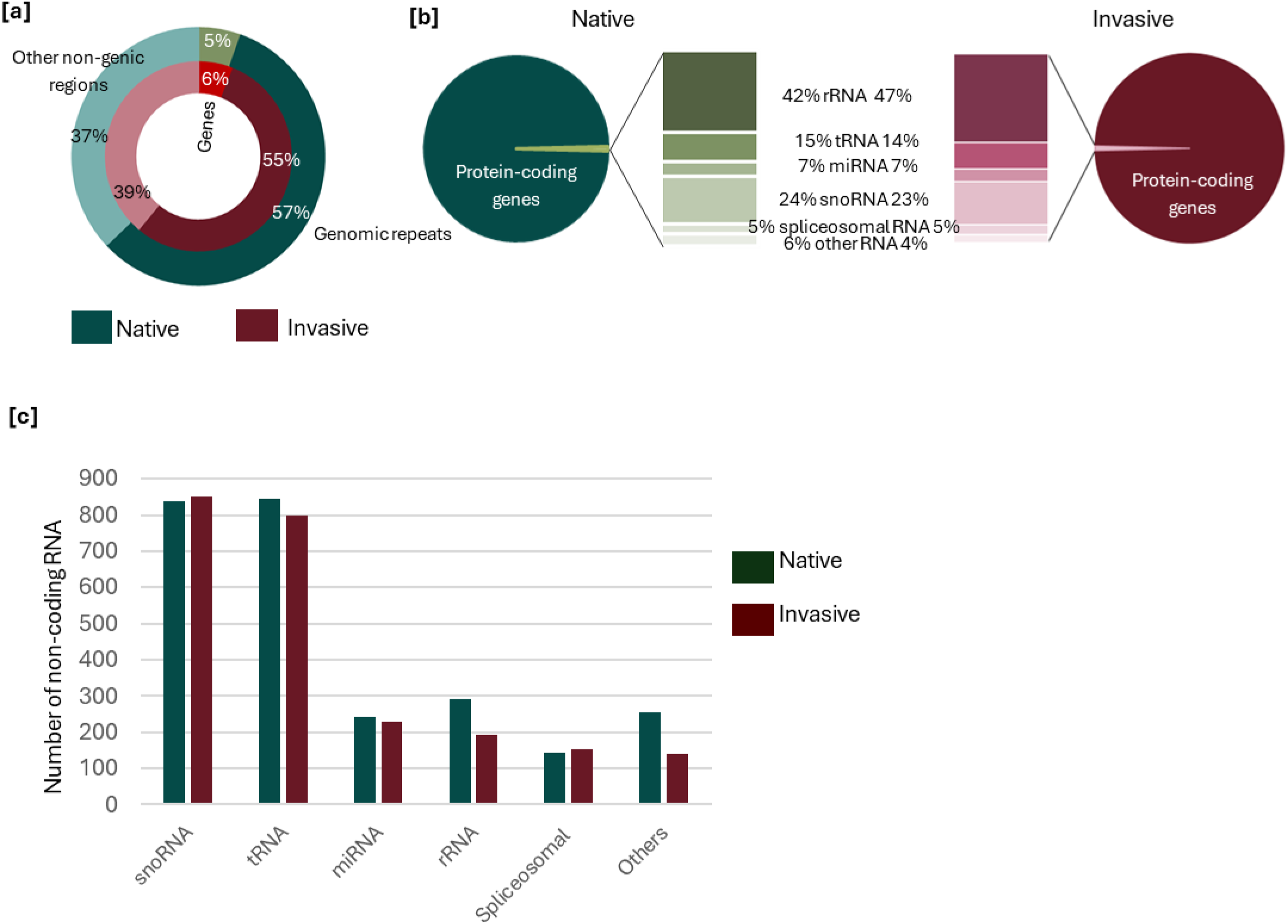
Gene composition of *P. australis* invasive and native genomes. [a] Genome composition based on length of genic and non-genic regions; [b] genome space assigned to different non-coding RNA classes compared to coding genes; [c] the annotated number of different non-coding RNA classes in each genome.

**Figure S5.**
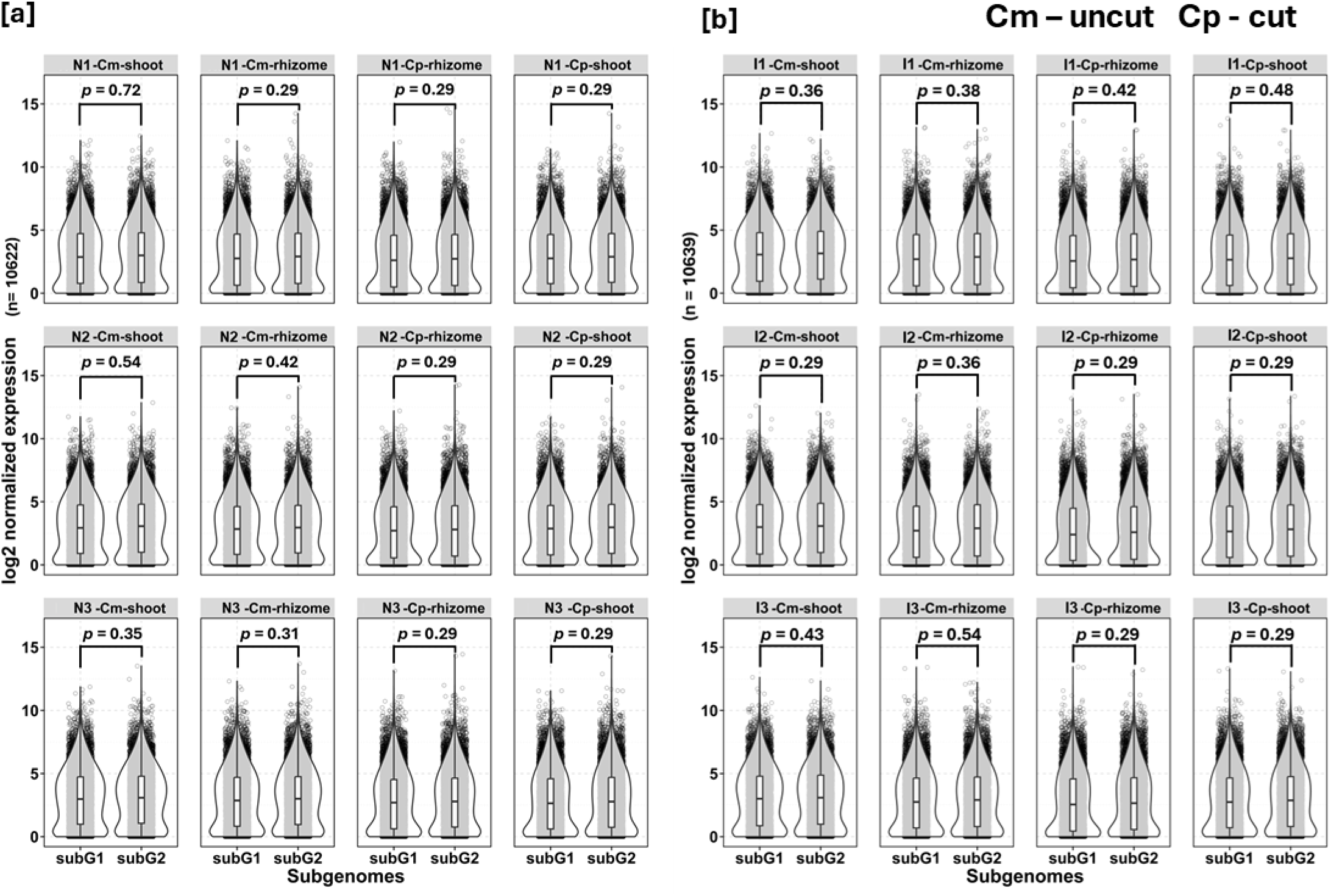
Homeolog expression across subgenomes. [a] native *P. australis* [b] invasive *P. australis.* Pairwise comparisons were performed for three genotypes in both shoot and rhizome tissues under cut and uncut conditions. P-values from paired Student’s t-tests, corrected using the Benjamini–Hochberg method, are shown for each comparison.

**Figure S6.**
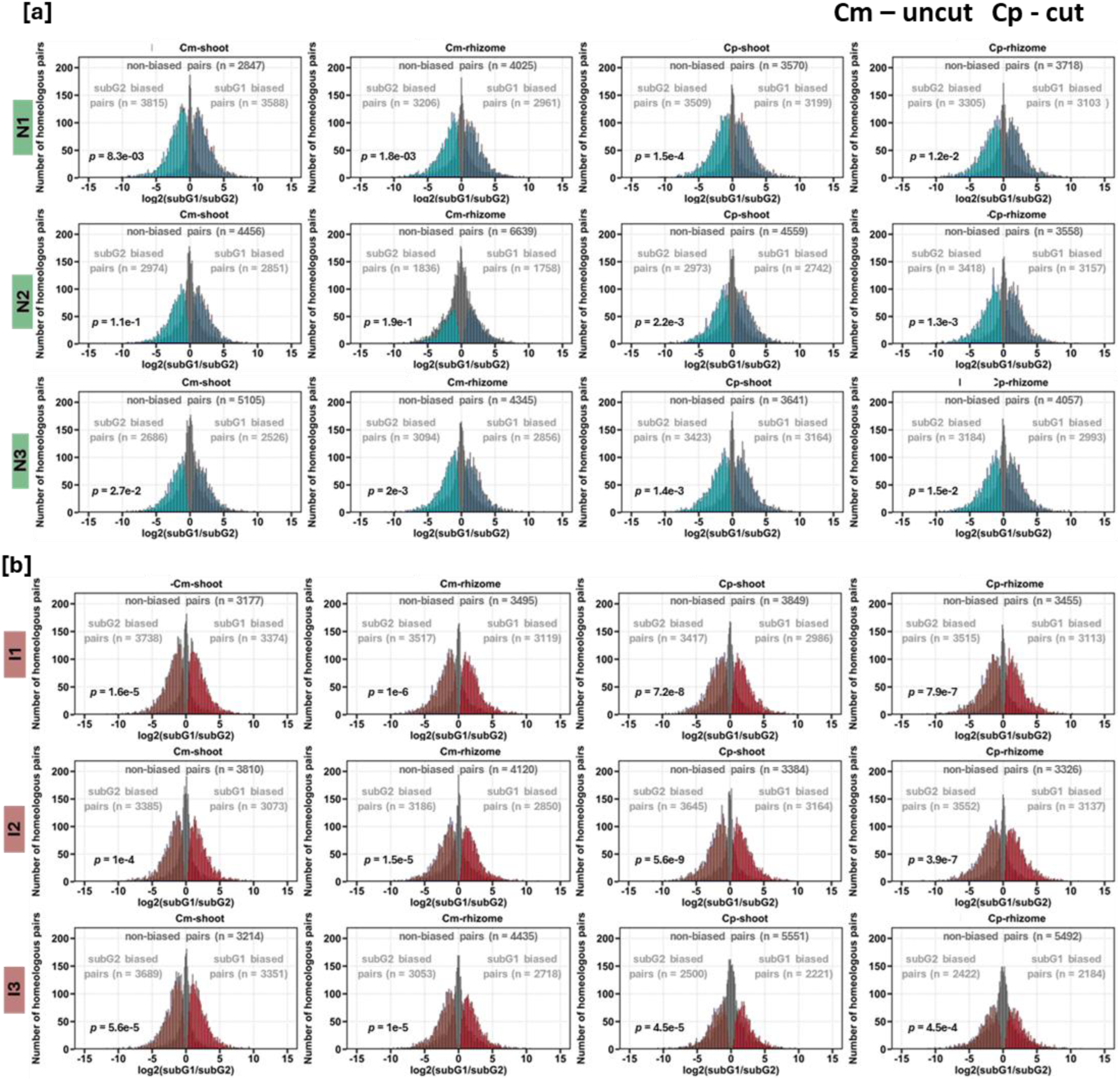
Homeolog expression bias in [a] native *P. australis* [b] invasive *P. australis.* Shoot and rhizome tissue from three genotypes under cut and uncut conditions were examined. Red regions indicate subgenome 1(subG1)-biased homeologs with log_2_ expression fold change > 0 and blue regions indicate subgenome 2(subG2) with log_2_ expression fold change < 0 at Benjamini-Hochberg adjusted *p*-value < 0.01 based on paired Student t-test. Chi-square test was used to determine whether the observed number of biased homeologs significantly deviates from equal sub-genome representation, and the Benjamini-Hochberg adjusted *p*-values are shown for each comparison.

**Figure S7.**
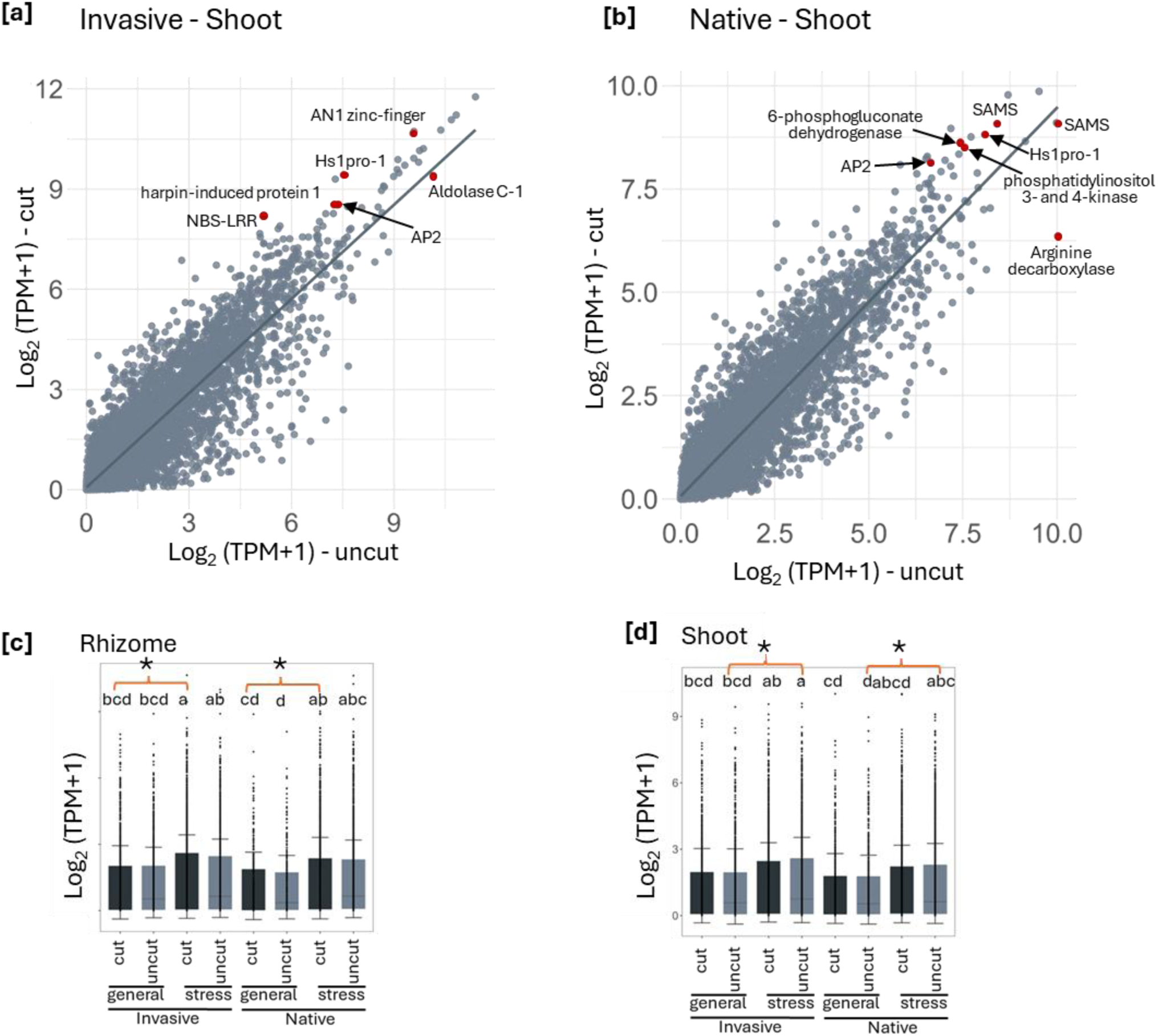
Expression of intronless genes and their association with stress responses in *P. australis*. Expression of intronless genes in [a] invasive and [b] native shoots. Red data points highlight highly expressed genes known for stress responsive functions in uncut and cut shoots. Expression of intronless genes categorized into stress and general responses in [c] rhizome and [d] shoot tissues. Genes were assigned to either stress or general metabolism based on GO annotations. Uncut shoots and cut rhizomes have more Significance assessed by one-way ANOVA (p < 0.05).

**Figure S8.**
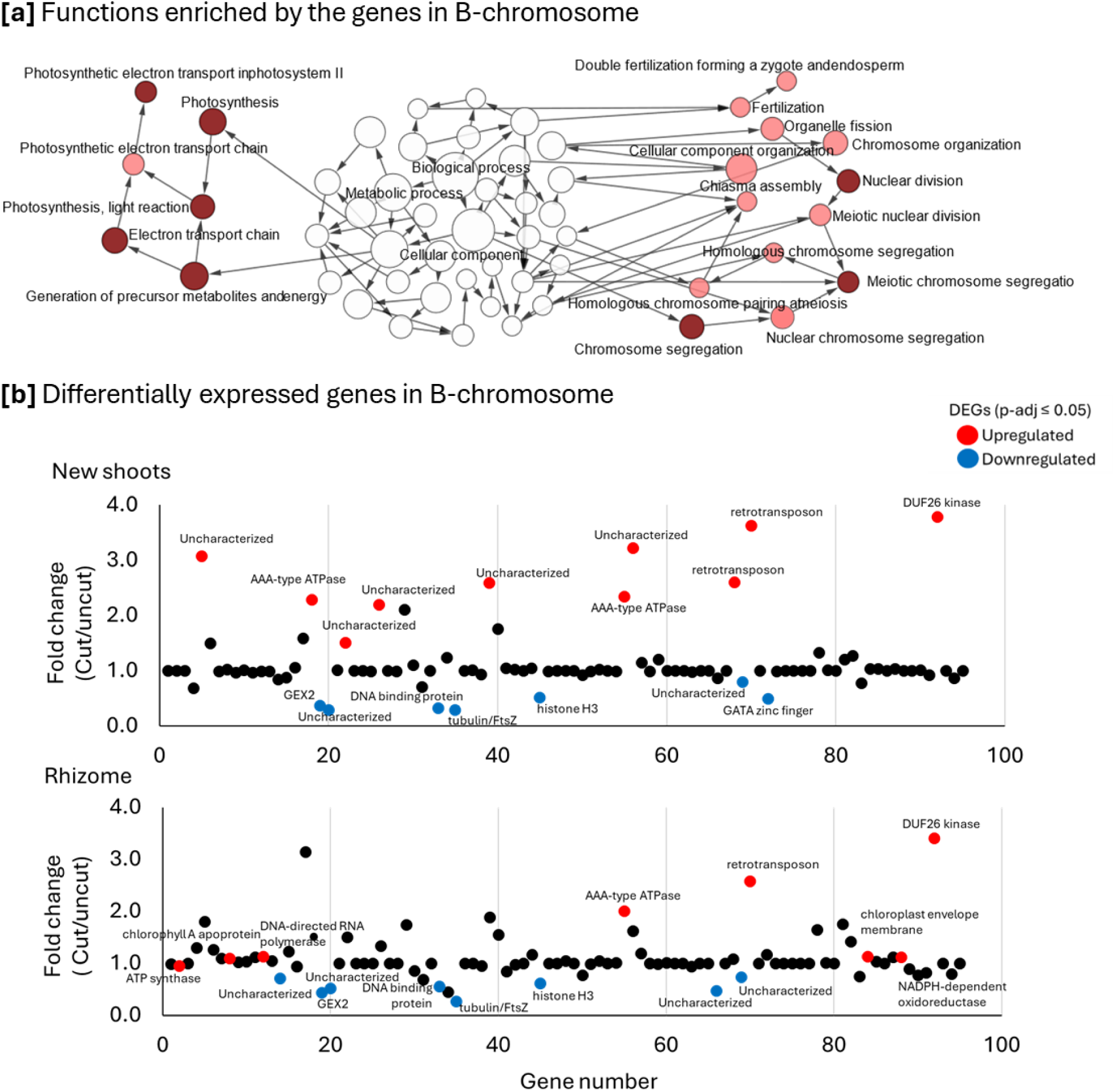
Gene functions and expression in response to cutback on the B chromosome. [a] Enriched functions represented by protein coding genes located in the B chromosome. The nodes in the functional cluster represent GO functions and the edges show sharedness of genes between functions. Two enriched functional clusters include photosynthesis and mitosis. [b] Differentially expressed genes (DEGs) in response to cutback. Ninety five genes were tested for DEGs at p-adj ≤ 0.5.

**Figure S9.**
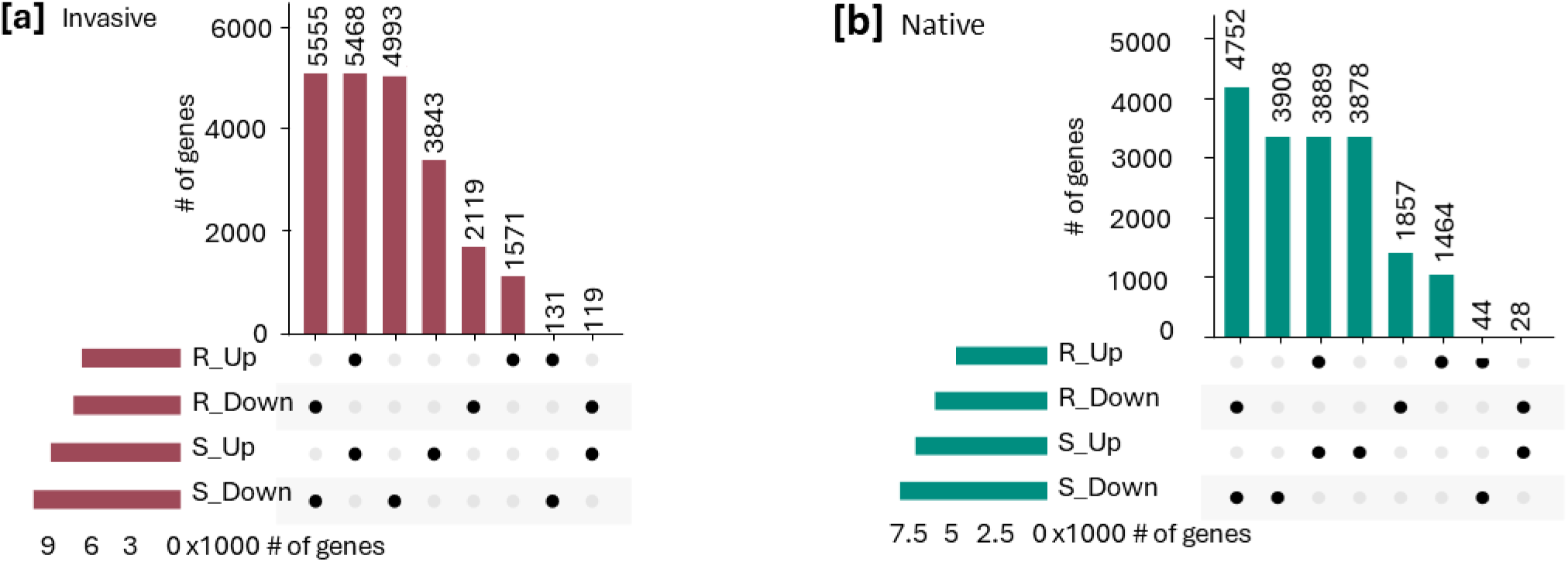
Differentially expressed genes (DEGs) induced or suppressed after cutback. [a] Invasive and [b] native subspecies independently tested for DEGs. DEGs between cut and uncut tissue were determined by p-adj values ≤ 0.05 based on Benjamini-Hochberg correction for multiple testing.

**Figure S10.**
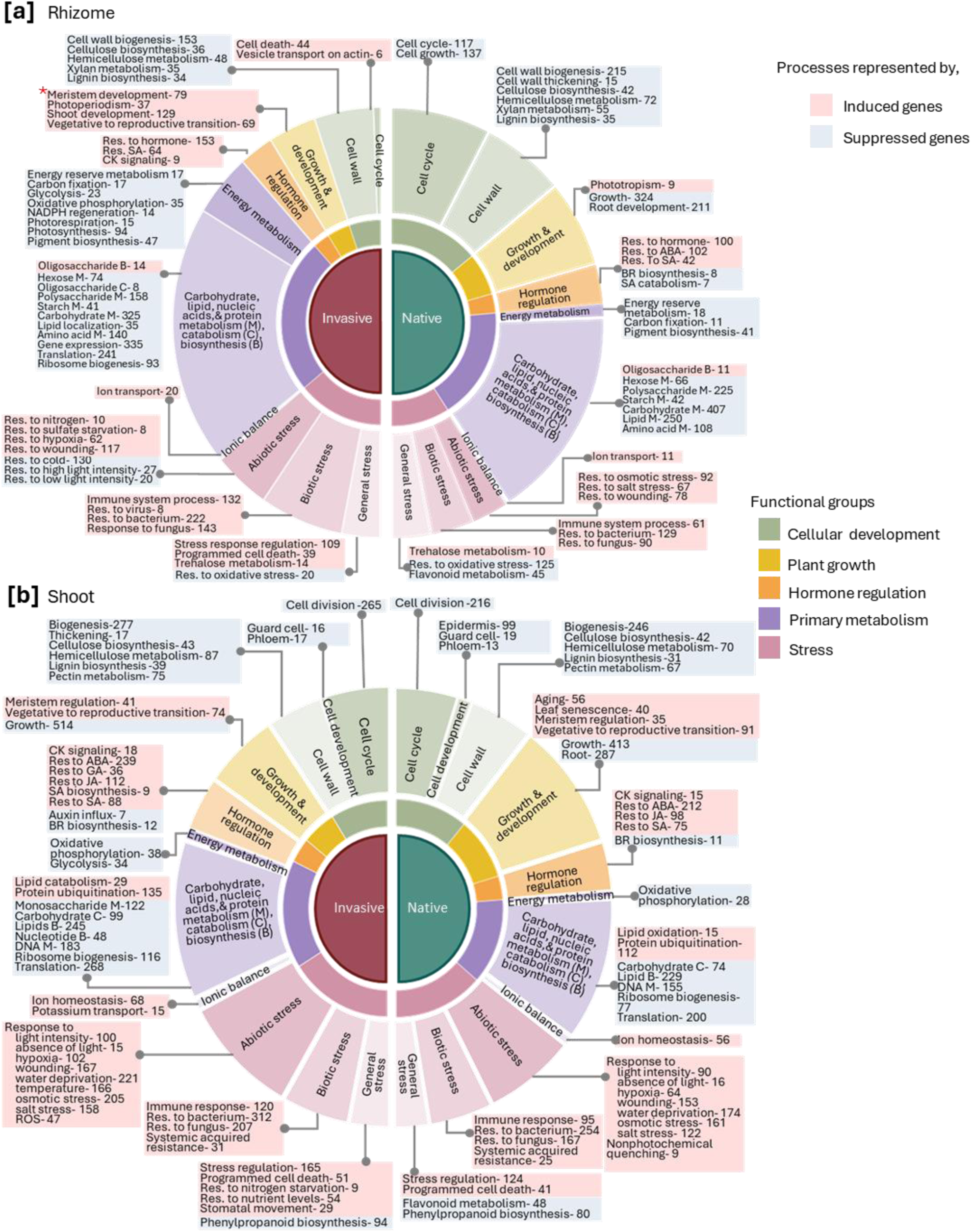
Summary of functionally enriched processes represented by DEGs. [a] Rhizomes and [b] shoots. Functional enrichment tested at adj p-values <0.05 with false discovery rate correction applied. The number of genes represented by a functional process are given in parenthesis within boxes under induced or suppressed genes.

**Figure S11.**
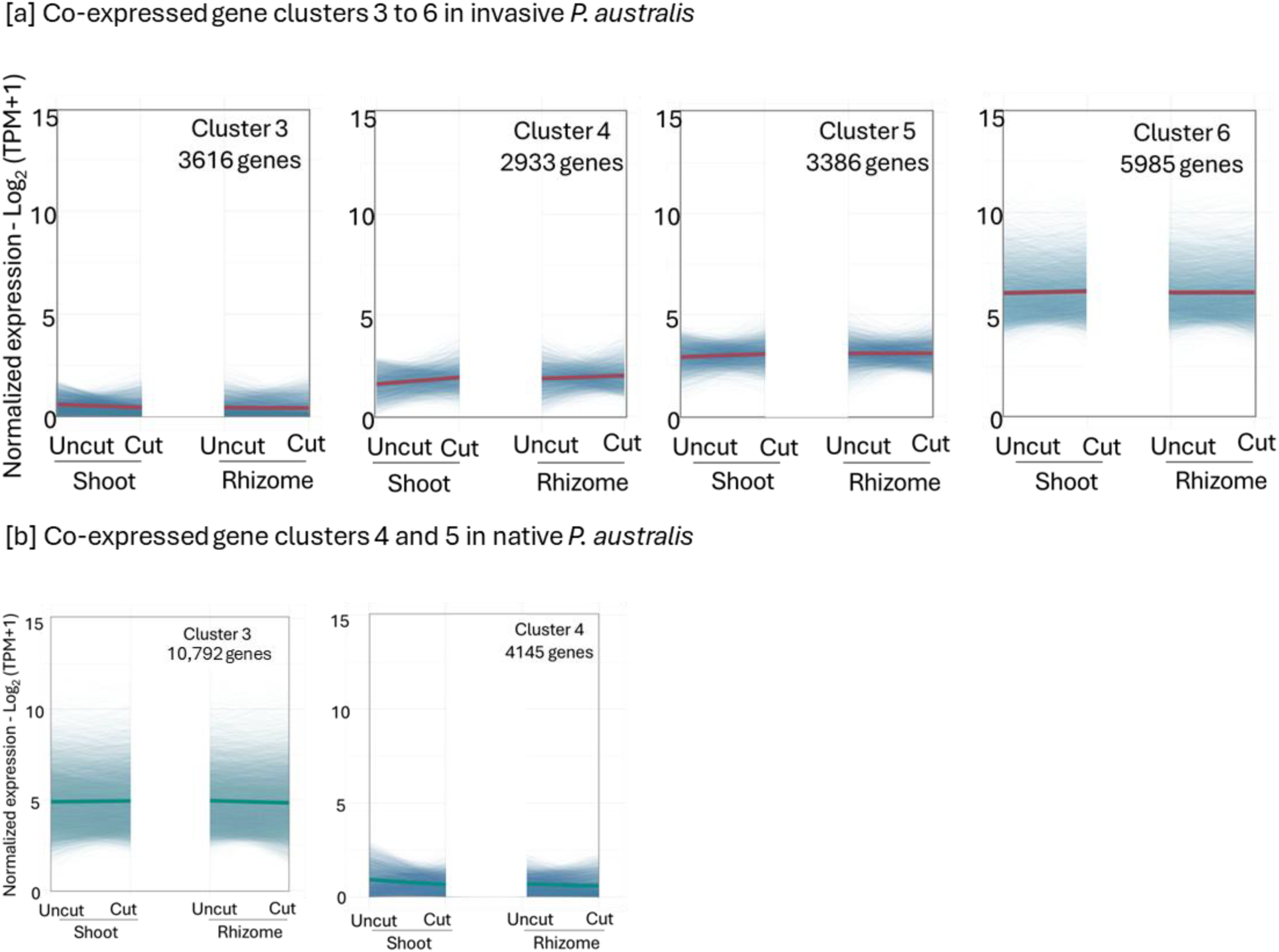
Co-expressed gene clusters in *P. australis* that are non-responsive to cutback in each tissue. Clusters 1-2 for each subspecies included in Fig. 5a and the remaining clusters shown here. Gene expression clusters based on co-expression profiles generated using k-means clustering performed with a membership cutoff of ≥ 0.5 across uncut and cut tissues from shoots and rhizomes of [a] invasive and [b] native subspecies. Thin lines represent individual genes contributing to the cluster while the thick line indicates the mean expression for the cluster in rhizome or shoots and uncut or cut samples.

**Figure S12.**
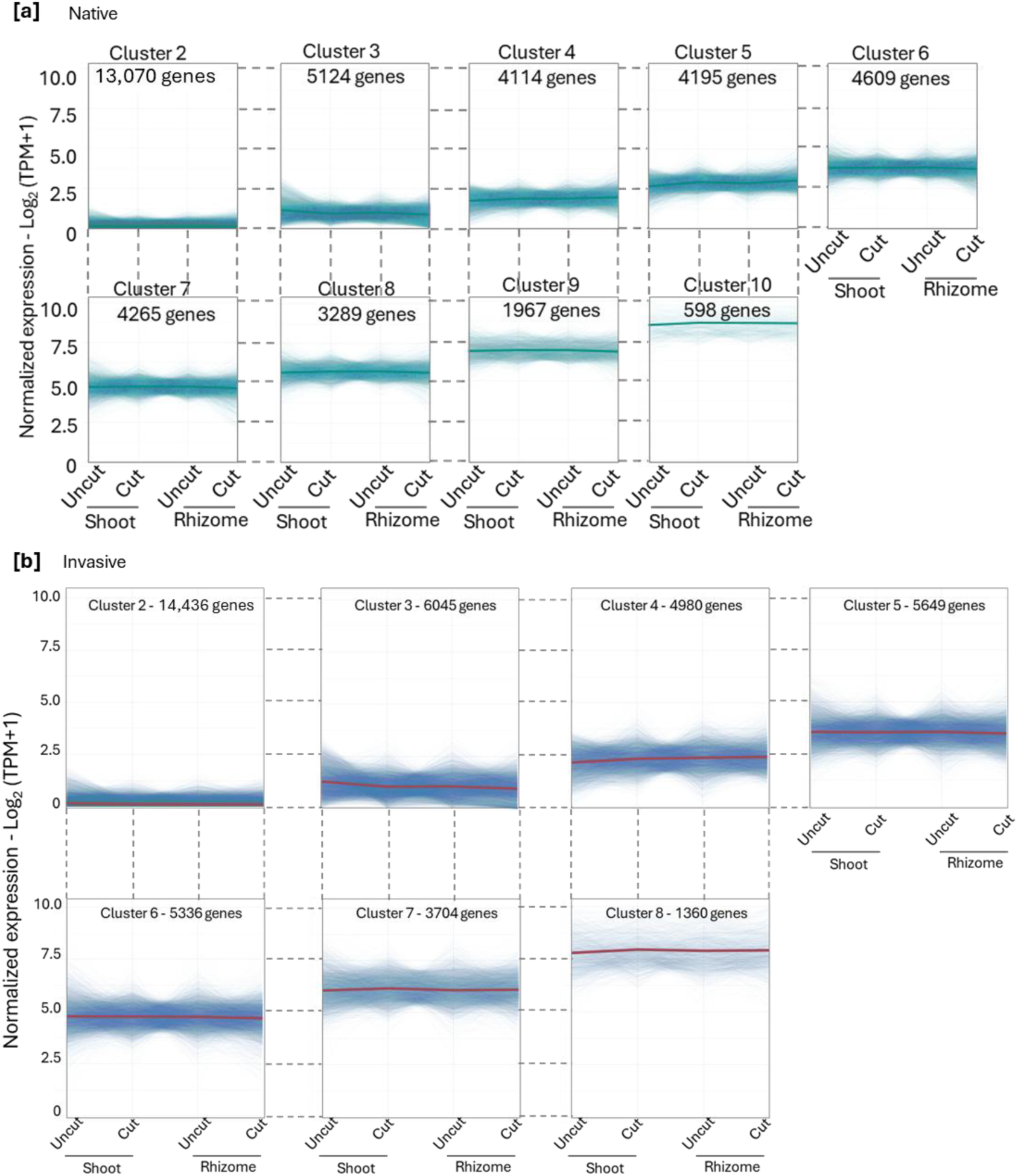
Clustering of all expressed genes coexpressed in [a] native and [b] invasive *P. australis.* Cluster 1 for each subspecies included in Fig. 5b and the remaining clusters shown here. k-means Clustering was performed with a membership cutoff of ≥ 0.5 across uncut and cut tissues from shoots and rhizomes, identifying co-expressed gene groups. Thin lines represent gene expression of individual genes in the cluster and the thick line represents mean expression.

**Figure S13.**
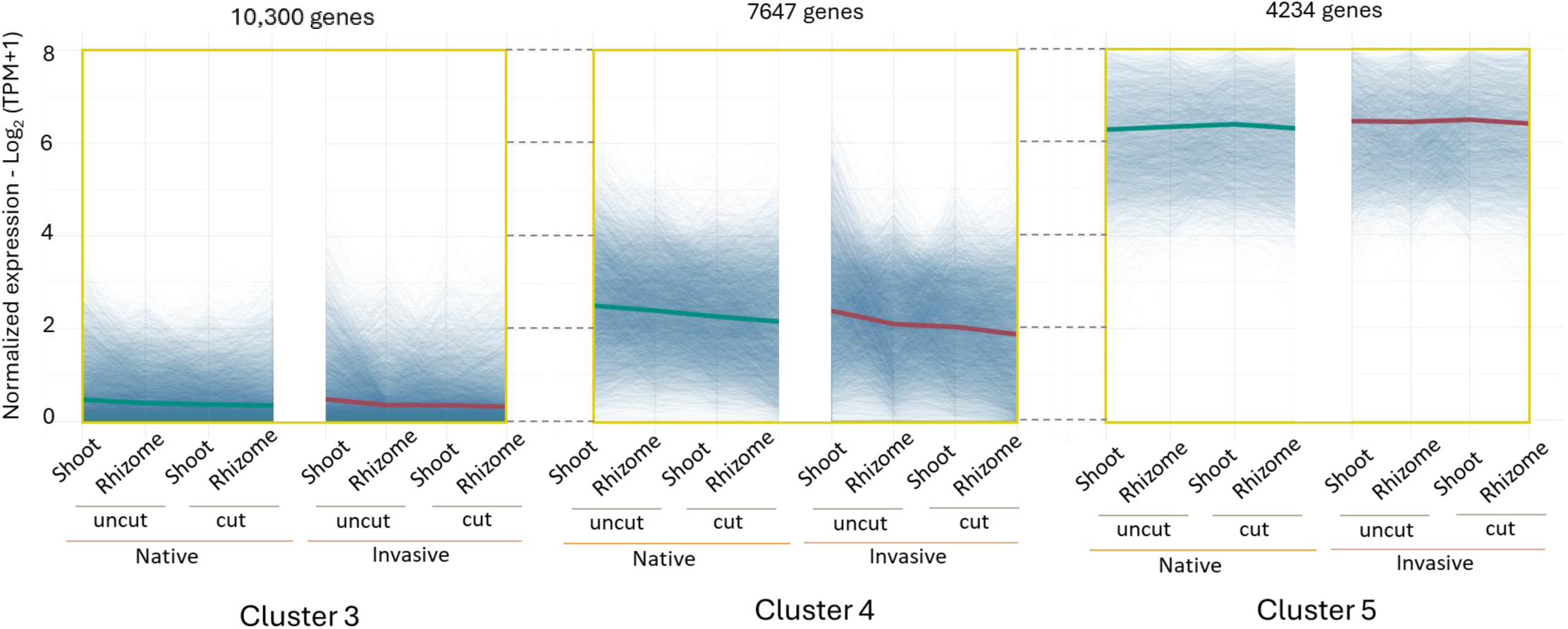
Co-expressed gene clusters of 1-1 orthologs between native and invasive *P. australis.* Clusters 3-5 shown here while the first two are included in Fig. 5c. k-means Clustering was performed with a membership cutoff of ≥ 0.5 across uncut and cut tissues from shoots and rhizomes, identifying similar co-expressed gene groups that capture transcriptomic responses to cutback. Thin lines represent gene expression of individual genes in the cluster and the thick line represents mean expression.

**Figure S14.**
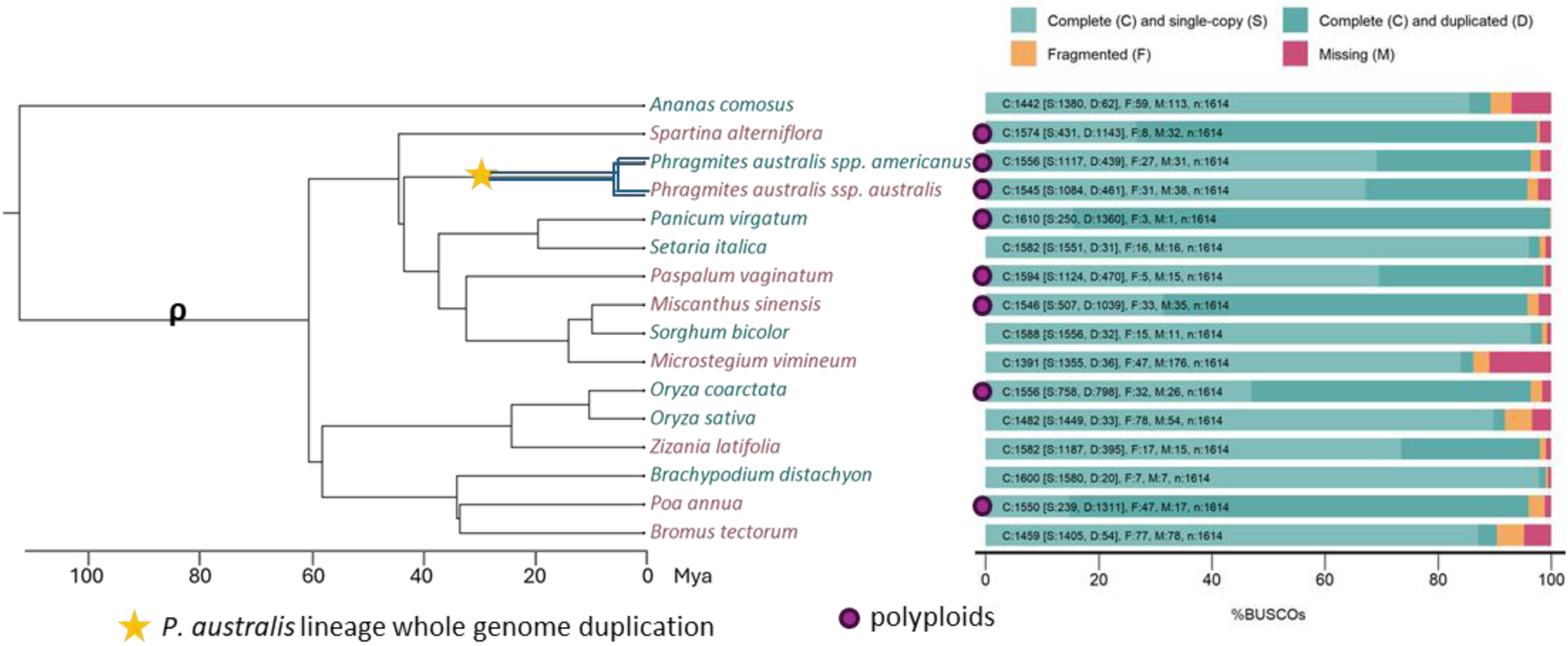
Genomes selected for invasive grass species and non-invasive relatives used as comparator species. Genome completeness assessed by BUSCO. Species reported to be invasive are in green text while non-invasive relatives are in magenta. Species with complete but duplicated copies (darker cyan) that exceed more than 5% of the complete single copy BUSCO genes (lighter cyan) are polyploids. An allopolyploidy event (∼30 Mya) followed by a lineage-specific genome duplication in *P. australis* is marked with a star; the ρ whole-genome duplication at the base of Poaceae is also indicated.

**Figure S15.**
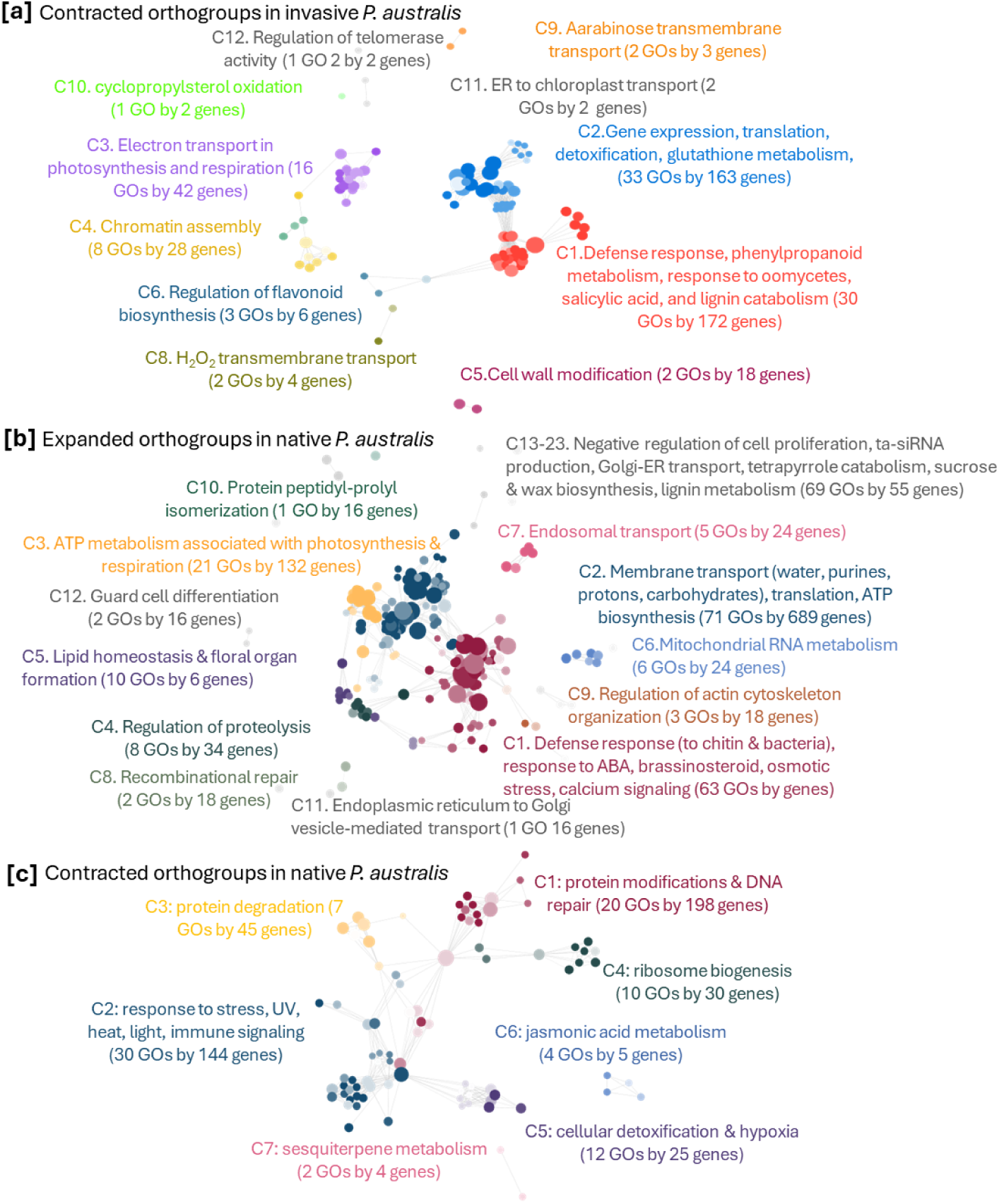
Enriched functions represented by *P. australis* orthogroups with altered copy number. [a] Contracted orthogroups in invasive, [b] expanded orthogroups in native, [c] contracted orthogroups in native *P. australis.* Nodes in clusters – gene ontology (GO) terms; node size – number of genes represented by that function; similar functions share the same color and are given a representative annotation; and the edges between nodes represent ortholog families that are shared between functions. All clusters included in the network have adj p-values <0.05 with false discovery rate correction applied. More significant values are represented by darker node colors.

**Figure S16.**
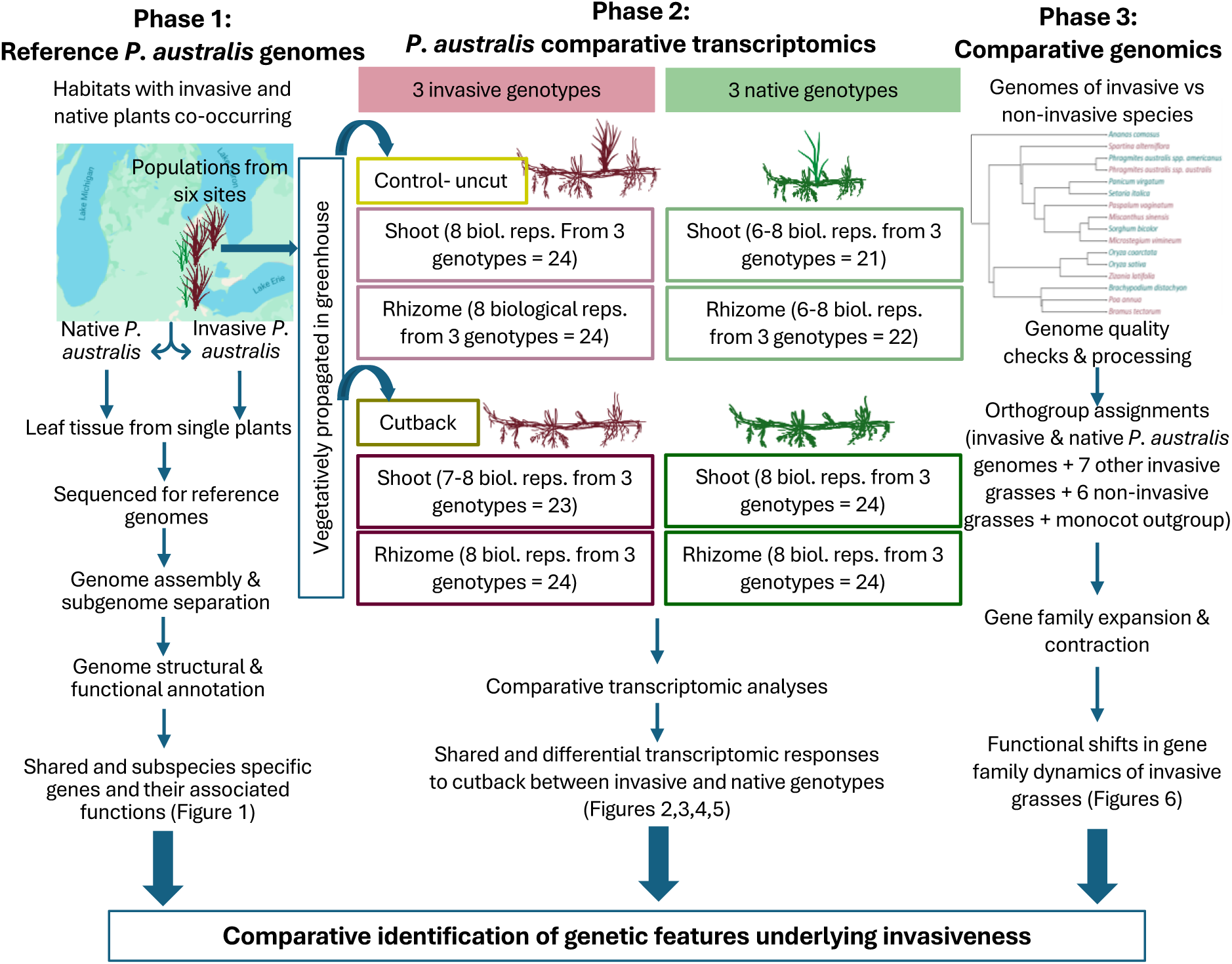
Experimental design and analysis overview. Phase I: Chromosome-level assemblies for invasive and native *Phragmites australis*. Phase II: Transcriptomic responses to cutback. Phase III: Comparative analyses across grasses to identify gene family expansions and contractions associated with invasiveness and their functions.

## Supplementary Tables

**Table S1**. Transposable elements annotated in the invasive and native *P. australis* genomes

**Table S2**. Protein coding gene model annotations based on rice homologs in the invasive (2.1) and native (2.2) *P. australis* genomes

**Table S3**. Ortholog expression and enriched functions. Ortholog pairs (3.1) and functions associated with 1:1 ortholog pairs conserved between the invasive and native *P. australis* genomes (3.2), enriched functions in single-copy orthologs (3.3), enriched functions of genes preferentially highly expressed in one subspecies (3.4), and functions enriched by genes highly expressed in one subgenome over the other in shoots and rhizomes of each subspecies (3.5)

**Table S4**. Transcriptomes sampled (4.1) and their normalized expression (TPM) in the invasive (4.2) and native (4.3) *P. australis* genomes

**Table S5**. Expression of genes in the B chromosome in response to cutback (5.1) and the enriched functions associated with the annotated genes (5.2)

**Table S6**. Differentially expressed genes (DEGs) in invasive shoots (6.1), invasive rhizomes (6.2), native shoots (6.3), and native rhizomes (6.4) in response to cutback and enriched functions represented by DEGs in shoots (6.5) and rhizomes (6.6)

**Table S7**. Genes in coexpressed clusters using DEGs (7.1), all genes (7.2), 1:1 ortholog pairs (7.3) clustered in invasive and native subspecies

**Table S8**. Genomes of invasive grasses compared to non-invasive relatives (8.1), their orthogroup gene assignment (8.2), the gene copy numbers assigned to each orthogroup (8.3), enriched functions associated with orthogroups expanded and contracted between *P. australis* subspecies (8.4), and GO functions enriched in orthogroups from grass species varying in copy number between invasive and non-invasive species (8.5). These functions are represented with the number of genes expanded or contracted in invasive and native *P. asutralis* as a focal pair of invasive-native species

## Supplementary Data Files

**File S1.** Genome sequence in fasta format for the native *P. australis ssp. americanus* reference genome

**File S2.** Genome sequence in fasta format for the invasive *P. australis ssp. australis* reference genome

**File S3.** Genome structural annotation in the GFF format for the native *P. australis ssp. americanus* reference genome

**File S4.** Genome structural annotation in the GFF format for the invasive *P. australis ssp. australis* reference genome

**File S5.** Scaffold order and assembled sequence of the B chromosome in the invasive *P. australis* genome

**File S6.** Protein coding sequence (CDS) file in fasta format for the native *P. australis ssp. americanus* reference genome

**File S7.** Protein coding sequence (CDS) file in fasta format for the invasive *P. australis ssp. australis* reference genome

## Data availability

All raw and assembled sequence data have been deposited to NCBI under BioProject PRJNA705976. The reference genome sequence, gene models, and RNA-Seq data are available at author (date; https://doi.org/10.5066/P13TOX3J).

## Author contribution

MD, KC, and KK conceived this research. MD directed the comparative genome and transcriptome analyses. KC, WB, and KK directed and performed field collections and greenhouse physiological experiments. DHO, PP, SW, and RG conducted genome assembly and annotation. SW, RG, TN, GW, PP and MD conducted data analysis. MD drafted the manuscript. The final version of the manuscript was generated with input from all authors. All authors approved the final version.

### Acknowledgements

The authors would like to thank The U.S. Fish and Wildlife Service and Michigan Department of Environment, Great Lakes and Energy for access to study sites and the Great Lakes Restoration Initiative for financial support. We thank Quynh Quach and Paul Richardson (Research assistants at Tulane University) for their assistance in coordinating group meetings, genotyping and maintenance of greenhouse stocks, and preliminary growth assays. This research was supported by the United States Geological Survey Cooperative Agreement G23AC00546. MD acknowledges the support of National Science Foundation awards NSF-IOS-EDGE-1923589 and DOE BER DE-SC0022985 for graduate student training. The authors also acknowledge the LSU High Performance Computing services for providing computational resources needed for data analyses. During manuscript preparation, ChatGPT (GPT-5.3, OpenAI, 2026) was used to correct typographic errors and clarity of the text. All AI-generated suggestions were reviewed and revised further and approved by the authors to ensure accuracy and maintain the integrity of the scientific content.

